# Long-read isoform sequencing reveals survival-associated splicing in breast cancer

**DOI:** 10.1101/2020.11.10.376996

**Authors:** Diogo F.T. Veiga, Alex Nesta, Yuqi Zhao, Anne Deslattes Mays, Richie Huynh, Robert Rossi, Te-Chia Wu, Karolina Palucka, Olga Anczukow, Christine R. Beck, Jacques Banchereau

**Author notes:** These authors contributed equally (Diogo F.T. Veiga, Alex Nesta). Correspondence (O.A.), (C.R.B.), (J.B.).

## Abstract

Tumors display widespread transcriptome alterations, but the full repertoire of isoform-level alternative splicing in cancer is not known. We developed a long-read RNA sequencing and analytical platform that identifies and annotates full-length isoforms, and infers tumor-specific splicing events. Application of this platform to breast cancer samples vastly expands the known isoform landscape of breast cancer, identifying thousands of previously unannotated isoforms of which ~30% impact protein coding exons and are predicted to alter protein localization and function, including of the breast cancer-associated genes *ESR1* and *ERBB2*. We performed extensive cross-validation with -omics data sets to support transcription and translation of novel isoforms. We identified 3,059 breast tumor-specific splicing events, including 35 that are significantly associated with patient survival. Together, our results demonstrate the complexity, cancer subtype-specificity, and clinical relevance of novel isoforms in breast cancer that are only annotatable by LR-seq, and provide a rich resource of immuno-oncology therapeutic targets.

## INTRODUCTION

Transcriptomic and proteomic diversity are influenced by alternative splicing (AS), transcription initiation, and polyadenylation in healthy and diseased cells (Baralle and Giudice, 2017; Manning and Cooper, 2017; Liu et al., 2017). Human tumors, including breast cancers, exhibit widespread changes in the AS isoform repertoire (Venables et al., 2008; Lapuk et al., 2010; Eswaran et al., 2013; Zhao et al., 2016), caused either by somatic mutation or mis-expression of the splicing regulatory machinery (Karni et al., 2007; Dvinge et al., 2016; Urbanski et al., 2018; Park et al., 2019). Specific spliced isoforms are important for cancer initiation, progression, metastasis, and drug resistance, with some AS events significantly linked to patient survival (Dvinge et al., 2016 Stricker et al., 2017; Urbanski et al., 2018; Read and Natrajan, 2018; Zhang et al., 2019b). For example, splicing of *CD44*, a transmembrane glycoprotein that functions in cell division, viability, and adhesion, has been linked with tumor progression and epithelial-to-mesenchymal transition in breast and ovarian cancer models (Brown et al., 2011; Olsson et al., 2011; Bhattacharya et al., 2018; Zhang et al., 2019a). Although the effects of a handful of spliced isoforms in cancer have been studied (Urbanski et al., 2018; Mitra et al., 2020), the clinical relevance of most isoform switches in tumors remain poorly characterized.

Global analyses of cancer transcriptomes have catalogued AS profiles in oncogenesis using short-read RNA-sequencing (RNA-seq) data, and have identified a number of recurrent and tumor-specific splicing alterations across many cancer types, including breast (Eswaran et al., 2013; Sebestyén et al., 2015; Zhao et al., 2016; Vitting-Seerup and Sandelin, 2017; Kahles et al., 2018). The detection and quantification of AS events using short-read RNA-seq data is inherently dependent on alignment of the RNA sequencing fragments to a reference genome and applying algorithmic reconstruction to identify cancer-associated isoforms. However, this approach often yields only a partial view of the splicing repertoire (Stark et al., 2019) because of limitations of transcript assembly tools. Indeed, current state-of-the-art spliced isoform reconstruction methods can only assemble ~20-40% of human transcriptomes (Steijger et al., 2013; Pertea et al., 2015; Song et al., 2019). Therefore, approaches that exclusively use short-read RNA-seq data are unable to fully characterize the cancer-associated AS isoform landscape, including the discovery of novel spliced isoforms involving non-adjacent exons.

Long-read mRNA sequencing (LR-seq) is able to accurately capture full-length isoforms from start to end, eliminating the need for reference-based transcript reconstruction (Sharon et al., 2013; Tilgner et al., 2014; Weirather et al., 2015; Nattestad et al., 2018; Lian et al., 2019). LR-seq of human and mouse cell and tissue transcriptomes has revealed a rich diversity of spliced isoforms (Tilgner et al., 2015; Lagarde et al., 2017; Byrne et al., 2017; Tilgner et al., 2018; Workman et al., 2019; Sheynkman et al., 2020). In cancer research, the use of LR-seq to identify primary tumor-associated spliced isoforms remains underexploited and has been limited to the study of human leukemia samples (Asnani et al., 2020; Tang et al., 2020). Importantly, the ability to aquire the depth of coverage needed to accurately quantitate transcripts using LR data is prohibitively expensive. Therefore, there is a need for a systematic application of LR-seq and subsequent analysis with short read RNA-seq to provide a more comprehensive view of the complexity of transcriptomes in primary tumors.

We use LR-seq and a multi-level analytical platform to thoroughly characterize the AS isoform landscape in breast cancer and normal breast samples. Our analyses identified tumor-specific isoforms, including isoforms associated with poor survival and specific breast cancer subtypes, and provide a library of novel breast tumor-specific isoforms as a resource for immuno-oncology therapeutic development.

## RESULTS

### LR-seq uncovers thousands of novel isoforms in human breast tumors

To interrogate the AS isoform landscape of breast cancer, we performed LR-seq on four normal human breast and 26 tumor samples. Our normal samples consisted of two cell lines and two primary tissues, and our breast cancer samples included 13 primary human breast tumor biopsies (three hormone positive - *ER*^+^/*PR*^+^, three *HER2*^+^, and seven triple-negative - TNBC), nine patient-derived xenograft (PDX) tumors, and four cancer cell lines (Fig. 1A, Table S1).

**Fig. 1.**
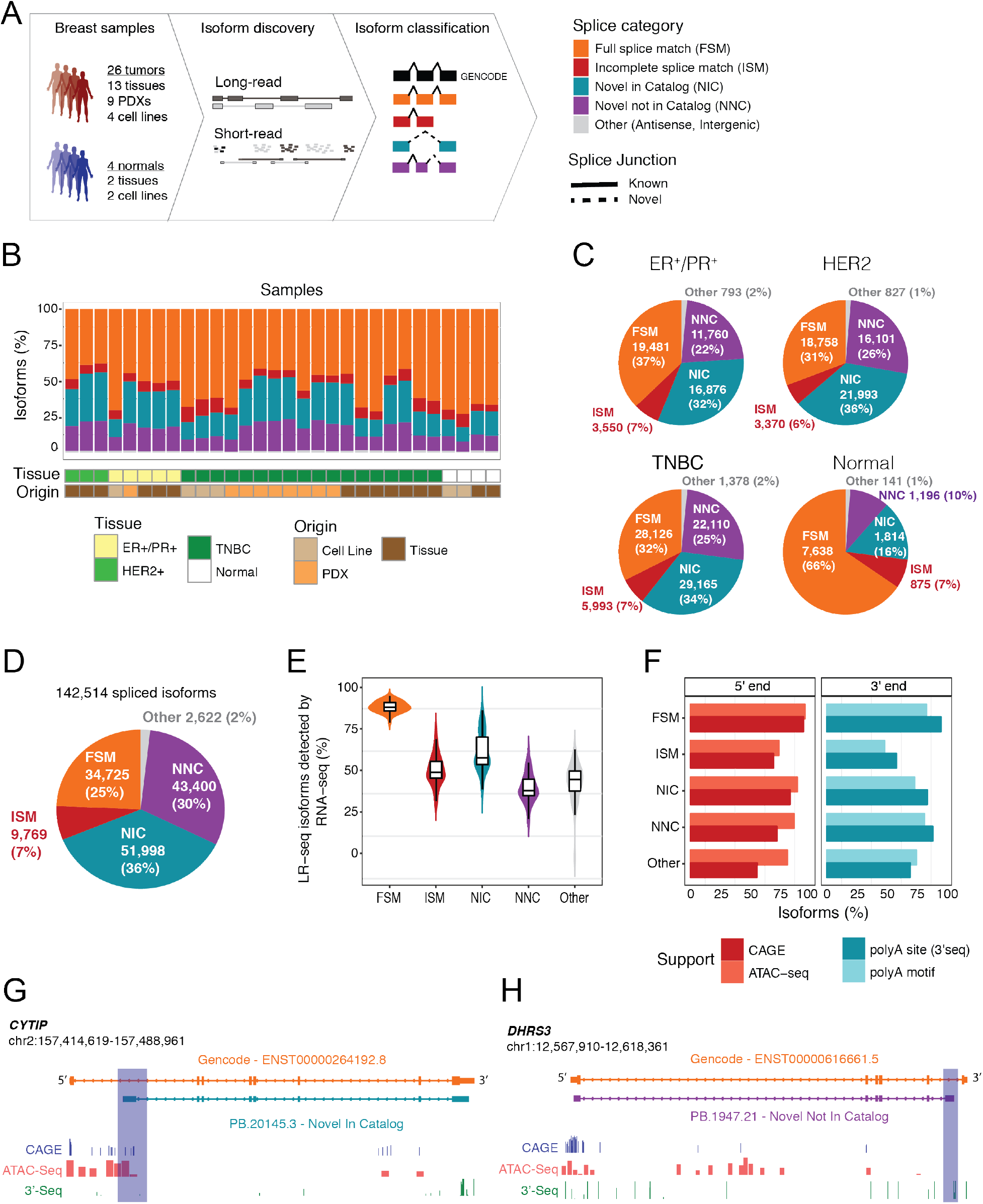
LR-seq identifies novel isoforms in breast cancer. **(A)** Schematic of isoform profiling by LR-seq and short-read RNA-seq and classification of breast cancer samples used in this study. LR-seq isoforms are classified based on their similarity to GENCODE v.30 isoforms using SQANTI isoform structural categories: FSM, ISM, NIC, NNC, and other (*e.g.*, antisense, intergenic, or readthrough transcripts), which are depicted schematically on the right. Novel splice junctions are depicted by dashed lines, and known junctions by solid lines. See also Fig. S1 and Table S1. **(B)** Classification of LR-seq isoforms detected in individual breast cancer or normal samples, colored by isoform structural categories from (A). Sample information including tissue type and origin are indicated below. See also Table S2. **(C)** Classification of isoforms obtained by LR-seq in samples from each breast cancer subtype (HER2^+^, ER^+^/PR^+^, TNBC) or normal. Isoforms are classified into structural categories as described in Fig. 1A. **(D)** Classification of LR-seq isoforms in the comprehensive LR-seq breast cancer transcriptome generated by merging all tumor and normal samples from (B). The percent and absolute number of distinct isoforms in each structural category from (A) are indicated. See also Fig. S2–3. **(E)** Percent of LR-seq isoforms detected by RNA-seq in 29 breast cancer and normal samples, plotted per isoform structural category from (A). (F) Percent of LR-seq isoform transcription start sites supported by CAGE (FANTOM5) or ATAC-seq (TCGA breast) peaks, or transcription termination sites supported by the presence of a poly(A) motif (SQANTI2) or 3’-seq peaks from the polyAsite database, plotted per isoform structural category from (A). **(G,H)** Structure of *CYTIP* (G) or *DHRS3* (H) novel NIC or NNC LR-seq isoforms compared to GENCODE isoforms, along with CAGE or ATAC-seq peaks supporting the novel transcription start site (G) and 3’-seq peaks supporting the novel transcription termination site (H). Novel regions are highlighted.

Isoforms obtained with Single-Molecule Real Time (SMRT) circular consensus sequencing (CCS) using the PacBio RSII and Sequel platforms were polished using the ToFU (Transcript isoforms: Full-length and Unassembled) pipeline (Methods, Fig. S1). A full-length isoform consists of a single mRNA molecule containing a poly(A) tail, where the entire transcript including cDNA adaptors at the 5’ and 3’ ends are successfully sequenced. After ToFU consensus clustering, 84% of CCS reads achieve 99.999% (Q50) accuracy (Table S2). Overall, per library, we obtained an average of 546,000 CCS reads, which after processing resulted in ~21,000 full-length polished isoforms (Table S2). As a quality control step after ToFU, we filtered transcripts with inadequate splice junction support and those that contained signatures of poly(A) intra-priming or non-canonical junctions derived from reverse transcriptase template switching (Methods, Fig. S1).

Next, isoforms were classified into known or novel isoforms based on their splice junction match to a reference transcriptome (GENCODE v.30) using SQANTI (Tardaguila et al., 2018). Known isoforms are classified as full-splice match (FSM), while novel isoforms include both transcripts that harbor a combination of known splice donors or acceptors that have not been previously cataloged in the same transcript (novel in catalog, NIC), as well as isoforms containing at least one novel splice site not present in GENCODE v.30 (novel not in catalog, NNC) (Fig. 1A). Overall, novel isoforms account for 17% to 55% of sequenced transcripts in the individual samples (average = 37%, Fig. 1B). Interestingly, the proportion of NIC and NNC isoforms is ~2-fold higher in all tumor subtypes versus normal samples (Fig. 1C). Finally, we construct an LR-seq breast cancer transcriptome by merging the 30 individual samples and removing redundant isoforms.

Our comprehensive LR-seq breast cancer transcriptome contains 142,514 unique full-length transcript isoforms (Fig. 1D) spanning 16,772 annotated genes and 905 novel loci, with a mean isoform length of 2.6 kb (Fig. S2A). Only a small fraction (2%) of poly(A) sequenced transcripts were novel antisense transcripts or mapped to intergenic regions (Fig. 1C). Two thirds of the breast cancer LR-seq isoforms (95,398 isoforms) were novel (NIC or NNC) (Fig. 1C), and mapped to regions of coding and non-coding genes. Within these NIC and NNC isoforms, LR-seq identified 67,727 unique splice junctions across 14,490 genes that were not previously annotated in GENCODE (Fig. S2B). The GC content adjacent to novel splice sites was higher than the known junction regions, suggesting that junctions in GC-rich regions may be under-represented when using traditional sequencing platforms (Fig. S2C). There was a positive correlation between the number of exons and number of novel LR-seq isoforms (Fig. S3), denoting that genes with higher exon complexity tend to generate a higher isoform repertoire. Finally, a large fraction of novel NIC (58%) and NNC (73%) isoforms were detected in only a single sample (Fig. S2D), while 19% of FSM isoforms are sample-specific. This may indicate that novel isoforms arise due to tumor heterogeneity as well as lack of coverage saturation in individual samples. Overall, breast cancer LR-seq identifies thousands of spliced isoforms which are not represented in current transcript databases.

### Novel breast cancer LR-seq isoforms are supported by orthogonal - omics data

To assess the support for LR-seq isoforms by short-read sequencing, we performed RNA-seq and quantified isoform expression in 29 out of our 30 LR-seq profiled breast samples. Briefly, 76bp-long paired-end RNA-seq libraries were sequenced at an average depth of 46 million reads per sample and mapped to our LR-seq breast cancer transcriptome using hisat2 and quantified using StringTie. While 89% of the annotated isoforms (FSM) were detected by RNA-seq (FPKM>0.5), novel NIC and NNC isoforms have average detection rates of 62% and 41% respectively (Fig. 1E).

In addition to RNA-seq, we used multiple orthogonal data sets to assess the reliability of novel breast cancer LR-seq isoforms, including CAGE (Cap Analysis Gene Expression), ATAC-seq (Assay for Transposase-Accessible Chromatin using sequencing) and 3’-seq. Novel 5’ isoform regions substantially overlapped with CAGE-validated transcription start sites (FANTOM5 CAGE) and open chromatin regions detected by ATAC-seq in TCGA breast cancer tumors (Corces et al., 2018) (Fig. 1F). Similarly, 3’ ends of novel LR-seq isoforms were supported by poly(A) motifs detected by SQANTI2 and *bona fide* transcription termination sites mapped using 3’-seq assays obtained from the polyAsite database (Fig. 1F). For example, our LR-seq breast cancer transcriptome identified a novel *CYTIP* isoform originating from an alternative transcription start site supported by proximal CAGE and ATAC-seq peaks (Fig. 1G). We also found a *DHRS3* isoform with a novel termination site supported by 3’-seq (Fig. 1H).

Altogether, the integration of LR-seq with orthogonal data reveals that ~80% of our novel (NIC and NNC) breast-cancer isoforms are validated by genomics (ATAC-seq) and/or transcriptomics (CAGE, 3’-seq) across independent samples.

### Breast cancer oncogenes and pathways are enriched in novel spliced isoforms

To assess the importance of novel isoforms from our LR-seq breast cancer transcriptome, we first examined the expression levels and gene pathways associated with these transcripts. Genes were binned into three groups based on our RNA-seq expression levels: low, average, and high based on FPKM cutoffs (Fig. 2A). Novel isoforms (NIC+NNC) were detected at similar rates for genes expressed at average and high levels (Fig. 2A), and at a lower rate for the lowest expressed genes, similar to FSM isoforms from GENCODE v.30. These data indicate that LR-seq detected novel transcripts even for lowly expressed genes, and that NIC and NNC isoforms from our LR-seq data are expressed at appreciable levels.

**Fig. 2.**
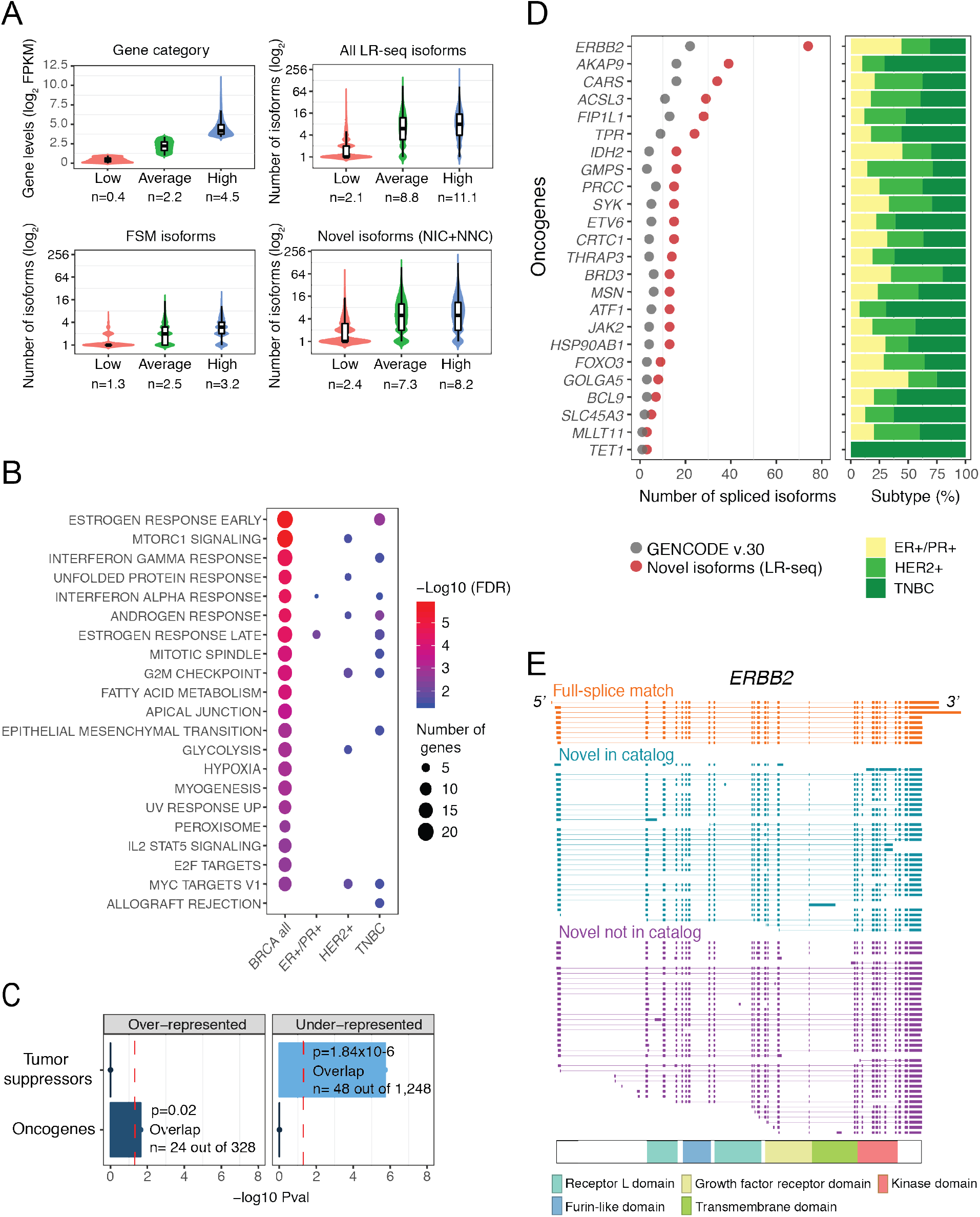
Novel LR-seq isoforms detected in breast tumors are enriched in cancer-associated pathways and oncogenes. **(A)** Correlation between gene expression levels from RNA-seq and number of transcript isoforms detected by LR-seq. Genes are binned based on quartile expression: low (1^st^ quartile), average (2^nd^ and 3^rd^ quartiles), and high (4^th^ quartile); where n is the mean log2 FPKM expression. Distribution of isoform numbers for each gene bin; where n is the mean absolute number of isoforms in the respective categories. **(B)** Pathways significantly enriched (MSigDB hallmark signatures; FDR<0.05) for genes with novel isoforms detected by LR-seq in all breast tumors or specific subtypes (HER2^+^, ER^+^/PR^+^, TNBC). Bubble size denotes the number of genes with novel isoforms in each pathway, and color denotes significance. See also Fig. S4A. **(C)** Enrichment analysis of oncogenes and tumor suppressors in genes with novel isoforms detected by LR-seq (hypergeometric test, *P*<0.05, cutoff indicated by a red dotted line). Oncogenes and tumor suppressor gene lists are obtained from MSigDB and TSGene databases, respectively. **(D)** Number of novel LR-seq isoforms compared to annotated GENCODE v.30 isoforms for selected oncogenes (left). The barplots (right) indicate the tumor subtypes where novel isoforms were detected, and are color according to subtypes described in Fig. 1B. **(E)** Structure of LR-seq *ERBB2* isoforms detected in breast tumors, grouped by isoform structural category from Fig. 1A. Included exons or introns are represented by solid boxes, spliced introns or exons by a line. The localization of *ERBB2* protein domains is indicated.

Next, we rank-ordered genes based on their ratio of isoform number gain when compared with GENCODE v.30 (#NIC+NNC isoforms / #GENCODE), and selected genes with >2-fold increase for pathway enrichment analysis. We performed this analysis for all combined breast tumors, as well individual breast cancer subtypes (Fig. 1B). Spliced genes with novel isoforms are strongly associated with key breast cancer pathways, including estrogen, androgen, and interferon gamma response, mTORC1 signaling, and mitotic spindle regulation (Fig. 2B, Fig. S4A). Other cancer relevant pathways are also over represented such as metabolism (glycolysis, hypoxia, fatty acid metabolism), replication (mitotic spindle, and G2M checkpoint pathways), and development (myogenesis, EMT). Myc targets were enriched in both HER2^+^ and TNBC tumors, while estrogen response was common to ER^+^/PR^+^ and TNBC. Notably, oncogenes are significantly over-represented when isoforms from all tumors are combined (Fig. 2C) while tumor suppressors are under-represented in this gene set (Fig. 2C).

We next examined individual genes that had a high gain of novel splice isoforms in our LR-seq breast cancer transcriptome. In total, 24 oncogenes including the human epidermal growth factor receptor 2 (*ERBB2*) exhibit a 2-fold increase in novel isoforms compared to GENCODE v.30 (Fig. 2D). *ERBB2* is often overexpressed in breast cancer due to gene amplification, and at least 3 spliced isoforms with clinical relevance have been identified (Castagnoli et al., 2019; Volpi et al., 2019). In addition to the 9 isoforms in GENCODE v.30, we detected 36 NIC and 38 NNC distinct spliced isoforms, revealing the complexity of *ERBB2* splicing regulation in breast tumors (Fig. 2E). Many of the *ERBB2* novel isoforms alter splicing of exons encoding known protein domains. We also found multiple novel spliced isoforms of genes significantly mutated in breast cancer, including *NCOR1, GATA3*, *SPEN*, and *PTEN* (Fig. S4B), as well as genes known to be alternatively spliced in cancer such as *CASP8*, *ENAH, BCL2L1*, and *STAT3* (Fig. S4C). In summary, LR-seq profiling of breast tumors identifies novel spliced isoforms in genes previously associated with key cancer pathways, as well as in known breast cancer oncogenes.

### Novel breast cancer LR-seq isoforms lead to alternative protein products

To understand the potential functional consequences of novel isoforms from our LR-seq breast cancer transcriptome at the protein level, we extracted Open Reading Frames (ORFs, *i.e.*, coding sequences) and predicted domains, transmembrane regions and subcellular localization using our ORF annotation pipeline, which includes Transdecoder for ORF predictions, as well as DeepLoc, TMHMM, and hmmer for localization predictions, and in-house scripts for comparative sequence analysis and nonsense-mediated decay (NMD) predictions (Methods, Fig. S1).

Overall, isoforms from all categories had very high coding potential (94-97%) based on our ORF prediction, except for antisense and intergenic transcripts for which 74% have a predicted ORF (Fig. S5A). However, NIC and NNC spliced isoforms are more likely to be targeted for mRNA degradation by the NMD pathway, since 11% of NIC and 20% of NNC translated ORFs contain premature termination codons compared to 3% of FSM ORFs (Fig. S5B). Similarly, novel ORFs absent from the protein coding database UniProt are subject to NMD at equivalent rates (11% and 22% of NIC and NNC, respectively) (Fig. S5B).

To determine whether LR-seq isoforms encode novel protein sequences, we compared the ORF of an LR-seq isoform to its closest match in UniProt using global pairwise alignment. The majority of annotated FSM (79%) and incomplete splice match (ISM) (85%) LR-seq isoforms encode ORFs that are >99% identical to an entry in UniProt (Fig. 3A). In contrast, only 23% of NIC or NNC LR-seq isoforms are annotated in UniProt (Fig. 3A). Thus, novel LR-seq isoforms are potential sources of novel proteomic diversity in breast cancer. We then investigated whether AS in our LR-seq breast cancer transcriptome leads to novel ORFs harboring changes in annotated protein domains, transmembrane regions or cellular localization. We found that ~20-30% of the novel ORFs lead to the loss of a transmembrane region or domain from the PFAM database (Fig. 3B), suggesting major changes in protein function or localization. In parallel, we used DeepLoc, a deep neural network-based tool (Almagro Armenteros et al., 2017), to predict the most likely subcellular compartment of LR-seq isoform-derived ORFs. We predicted that a third of the novel protein isoforms (25,714 ORFs) would change their subcellular localization compartment compared to their corresponding canonical UniProt entry (Fig. 3C). The localization switches are found primarily between cytoplasmic and nuclear localized protein isoforms (7,580 ORFs), followed by cytoplasmic and mitochondrial changes (3,777 ORFs) (Fig. S5C). Therefore, AS in breast cancer often leads to changes in protein localization which might impact spliced isoform function.

**Fig. 3.**
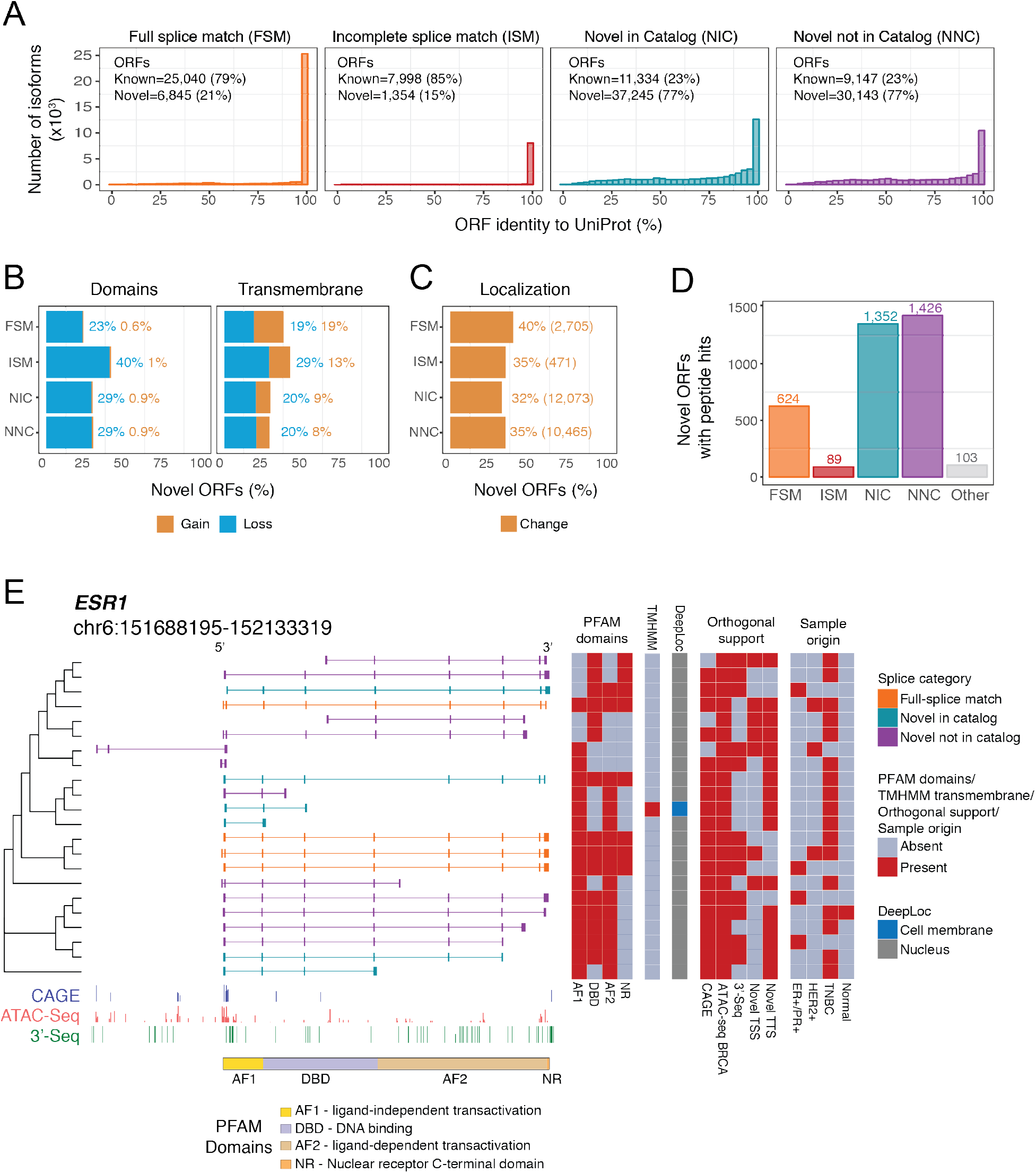
Novel LR-seq isoforms detected in breast tumors are predicted to impact protein sequence, domains, or localization. **(A)** Percent of amino acid sequence identity for LR-seq isoforms-derived Open Reading Frames (ORFs) compared to their closest human protein isoform in UniProt, plotted by isoform structural category from Fig. 1A. Known ORFs exhibit > 99% identity and novel ORFs <99% identity with UniProt. See also Fig. S5. **(B)** Percent of novel LR-seq isoform-derived ORFs predicted to gain or lose a conserved PFAM domain or transmembrane region compared to their closest human protein isoform in UniProt. **(C)** Percent of novel LR-seq isoform-derived ORFs predicted by DeepLoc to exhibit a different subcellular localization compared to their closest human protein isoform in UniProt. The absolute number of ORFs in each structural category is indicated. See also Fig. S5C. **(D)** Number of novel LR-seq isoform-derived ORFs validated by MS/MS proteomics, plotted per isoform structural category from Fig. 1A. Peptide search was conducted using 275 breast cancer samples (170 patients) from Clinical Proteomic Tumor Analysis Consortium (CPTAC). **(E)** LR-seq *ESR1* isoforms detected in breast tumors and predicted changes in protein domains and localization. LR-seq isoforms are grouped based on ORF similarity. For each isoform, the presence of PFAM domains, shown schematically at the bottom (AF1 – ligand-independent transactivation domain; DBD – DNA binding domain; AF2 – ligand-dependent transactivation domain; NR – Nuclear receptor C-terminal domain), as well as transmembrane regions (TM) identified by TMHMM, and predicted subcellular localization (DeepLoc) are indicated in adjacent heatmaps. Supporting orthogonal data such as the presence or absence of CAGE or ATAC-seq peaks (ATAC-seq BRCA) supporting novel transcription start sites (TSS), or 3’-seq peaks supporting novel transcription termination sites (TTS) is indicated. The sample origin where the isoform is detected, including breast cancer subtype (ER^+^/PR^+^, HER2^+^, TNBC) is indicated. Genomic tracks for CAGE, ATAC-seq and 3’-seq along with the localization of *ESR1* protein domains are displayed at the bottom.

Beyond transcript annotation, our pipeline leverages existing proteomics data for isoform validation (Fig. 3D). To determine the rate of isoform detection by MS/MS proteomics, we performed *in silico* peptide identification using our LR-seq-derived ORFs. We then intersected our data by spectral matching between theoretical peptides derived from LR-seq ORFs and experimentally-mapped peptides from 275 publicly available breast tumors samples (170 distinct patients) profiled by MS/MS proteomics by the Clinical Proteomic Tumor Analysis Consortium (CPTAC), including 125 TCGA patients (Mertins et al., 2016) and an additional 45 patient cohort (Johansson et al., 2019). The proteomic analysis found isoform-specific peptides supporting 1,352 NIC and 1,426 NNC novel LR-seq-derived ORFs (Fig. 3D). Additionally, we also identified 624 annotated FSM isoforms producing novel ORFs not present in UniProt. Although comprehensive tandem mass spectrometry (MS/MS) validation of isoforms with unique peptides is challenging, as most peptides map to regions shared by multiple isoforms (data not shown), these data suggest that our predicted ORFs are both transcribed and translated.

In sum, our analytical pipeline reveals that novel spliced LR-seq isoforms detected in breast cancer encode novel protein isoforms with changes in functional domains, transmembrane regions and/or cellular localization, and that the translation of novel isoforms can be confirmed with MS/MS data.

### Novel *ESR1* spliced isoforms predicted to impact protein localization are expressed in breast tumors

We next applied our isoform annotation pipeline to investigate the predicted functional effects of alternative splicing in *ESR1* (ERα), a clinical biomarker of hormone-positive breast cancers with several AS isoforms associated with cancer progression or treatment (Thomas and Gustafsson, 2011). In total, we detected 22 protein-coding isoforms in the *ESR1* locus, with 18 NIC or NNC isoforms being absent from GENCODE v.30 (Fig.3E). Among those, 7 novel protein isoforms are predicted to lack the DNA binding domain. 11 *ESR1* isoforms contained with novel regions affected by AS, including 5 protein isoforms with loss of the ligand-independent transactivation domain (AF1) and 6 protein isoforms with loss of the ligand-dependent transactivation domain (AF2) (Fig.3E). A unique *ESR1* isoform was predicted by TMHMM to contain a transmembrane domain and by DeepLoc to be localized to the cell membrane (Fig. 3E). This isoform could be involved in therapy resistance, as ESR1 localization to the plasma membrane promotes agonist action and resistance to tamoxifen (Thomas and Gustafsson, 2011).

Overall, characterization of the *ESR1* isoform diversity indicates that our LR-seq breast cancer transcriptome, coupled with protein-level annotation, can identify AS isoforms with putative novel functions and possible clinical relevance in breast cancer.

### Tumor subpopulations can be clustered by distinct splicing signatures

To specifically identify LR-seq isoforms enriched in breast tumors *vs.* normal tissues, we analyzed AS events in 1,135 human breast tumors and 1,443 normal tissue samples from TCGA and GTEx. We used SUPPA2 (Trincado et al., 2018) to extract 310,861 AS events in these 2,579 RNA-seq samples (Fig. 4A), using isoforms unique to our LR-seq breast cancer transcriptome (NIC and NNC) and annotated GENCODE v.30 transcripts as a reference. SUPPA2 quantifies AS events using percent spliced-in (PSI) - which measures the ratio of isoforms harboring the AS event across seven types of events: skipped exon (SE), mutually exclusive exons (MX), alternative 5’ splice-site (A5SS), alternative 3’ splice-site (A3SS), retained intron (RI), alternative first exon (AF), and alternative last exon (AL).

**Fig. 4.**
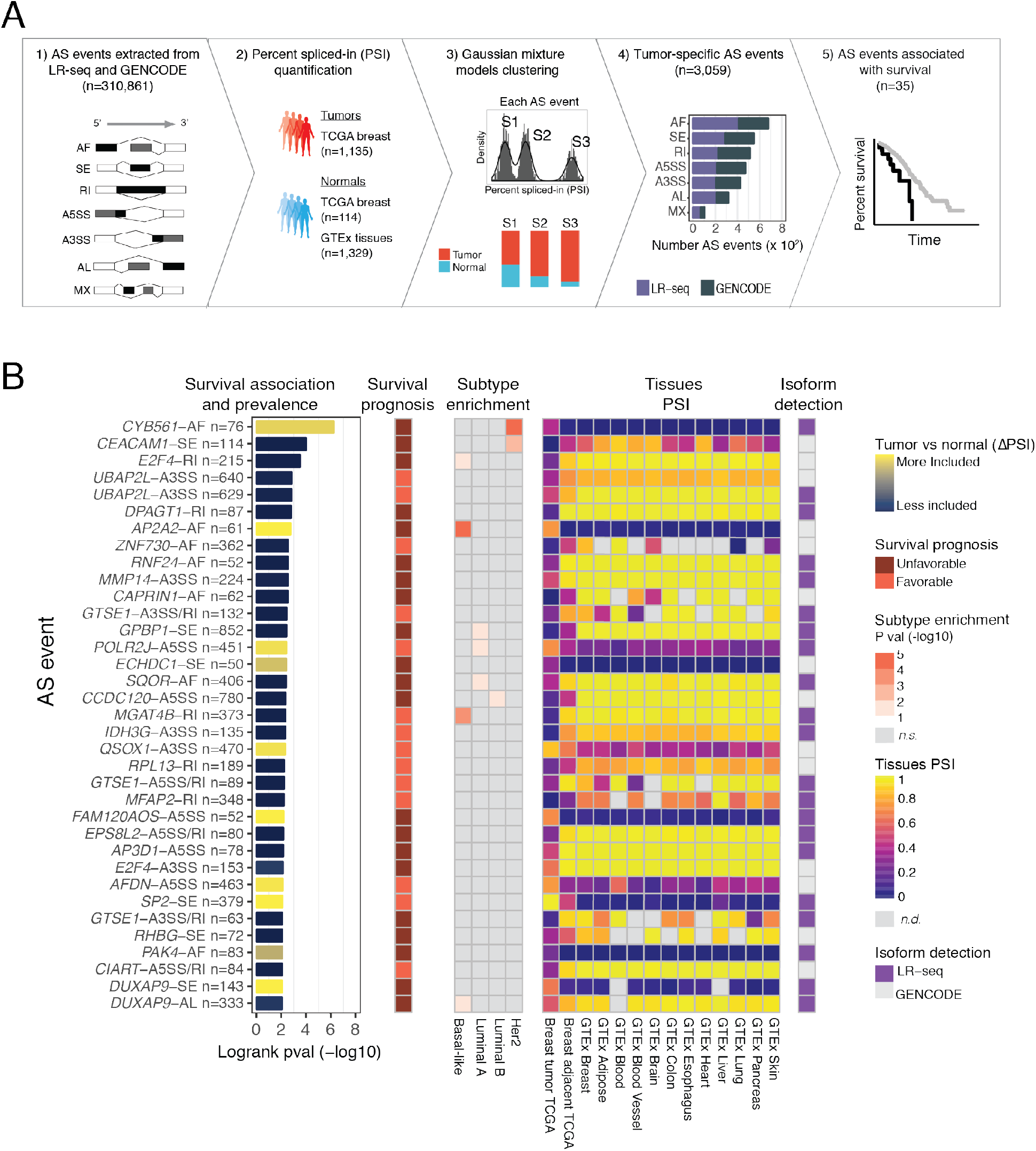
Patient clustering identifies splicing alterations associated with overall survival in breast cancer. **(A)** Identification of tumor-specific AS events in TCGA breast cancer patient subpopulations using the Gaussian Mixture Models (GMM) clustering approach. Seven types of AS events (SE, skipped exon; MX, mutually exclusive exons; A5SS, alternative 5’ splice-site; A3SS, alternative 3’ splice-site; RI, retained intron; AF, alternative first exon; AL, alternative last exon) are extracted from both LR-seq and GENCODE v.30 isoforms (1), and quantified, as percent-spliced-in (PSI) with SUPPA2 using RNA-seq from 2,579 samples including TCGA breast tumors and normal tissues from TCGA and GTEx (2). The GMM clustering approach provides for each AS event the optimal number of distinct sample subpopulations (*e.g.*, S1-S3) that fit the PSI distribution, as well as the frequency of tumor and control samples in each subpopulation (3). The GMM clustering approach identifies 3,059 tumor-specific AS events in TCGA breast tumors *vs.* normal tissues, plotted per AS event type (4). The Kaplan-Meier survival analysis compares survival rates in the identified subpopulations for all tumor-specific events, and detects 35 AS events associated with subpopulations with differential survival in TCGA (5). **(B)** Tumor-enriched AS events associated with overall survival in TCGA breast tumors identified by the GMM clustering approach from (A). Only AS events detected in ≥ 50 patients, with |ΔPSI | ≥ 20, and with significant survival association are shown, ranked by differential survival (logrank test, adj. *P*<0.01). AS events are labelled with gene name, AS event type, and number of patients, and colored based on inclusion levels (ΔPSI) in tumors *vs.* normal tissues. Related information for each AS event is depicted in heatmaps, including survival prognosis, breast tumor subtype enrichment, tissues PSI values, and source of isoform detection (*n.s.*, not significant; *n.d.*, not detected).

Given the heterogeneity of breast cancer, which can be classified into different subtypes based on gene expression and AS levels (Perou et al., 2000; Kahles et al., 2018), we introduced a novel approach for stratifying patients into groups based on distinct splicing alterations when compared to control samples. Our Gaussian Mixture Models (GMM) clustering approach simultaneously groups tumors and normal samples based on AS expression patterns, then identifies, for each splicing event, several clusters (*i.e.*, sample subpopulations) with two major features: (*i*) high frequency of tumor samples, and (*ii*) significant differential splicing (∆PSI) compared to normal tissues (Fig. 4A). Overall, our GMM clustering analysis identified 3,059 tumor-specific AS events in breast cancer with |∆PSI| ≥ 20% in subpopulations of at least 50 patients (Fig. 4A). Of those, 1,638 AS events (54%) were derived from isoforms present in our LR-seq breast cancer transcriptome and not annotated in GENCODE v.30, which highlights the contribution of novel isoforms in tumor-associated splicing (Fig. 4A). Therefore, our GMM clustering approach identified recurrent AS events in breast cancer, and found that they are often restricted to a subpopulation of patients in TCGA.

### Discovery of tumor-specific splicing events associated with survival

To distinguish isoforms associated with breast cancer prognosis, we directly compared the overall survival rates of each of the 3,059 tumor-specific AS events identified by GMM clustering. A total of 35 AS events in 30 distinct genes correlated with survival in TCGA (Fig. 4B, Table S3). The most highly associated AS events with a decrease in overall survival are an alternative first exon in *CYB561*, a skipped exon in *CEACAM1*, and loss of an intron retention event in *E2F4*. Genes containing AS events associated with overall survival are known components of cancer-related pathways, including regulation of transcription (*E2F4*, *ZNF730, GPBP1*, *POLR2J*, *SP2*, *CIART*), cell cycle (*E2F4*, *GTSE1*), or cell-cell adhesion (*CEACAM1*, *MMP14, EPS8L2, AP3D1*, *AFDN*, and *PAK4*) (Fig. 4B).

Ten of the 35 overall survival-associated AS events are enriched in specific breast cancer subtypes. For example, events in *CYB561* and *CEACAM1* are enriched in *HER2+*, and *E2F4*, *AP2A2, MGAT4B*, and *DUXAP9* events are enriched in basal-like breast tumors (Fig. 4B). Finally, 21 of our overall survival-associated AS events (~60%) were absent from GENCODE v.30, including the events in *CYB561*, *UBAP2L*, and *DUXAP9.* This indicates the importance of LR-seq in developing reference isoform transcriptomes that can elucidate clinically-relevant AS events.

Among the AS events associated with survival differences, we identified an exon skipping event in the cell adhesion molecule *CEACAM1* in 114 TCGA breast cancer patients (Fig. 4B, Fig. 5A-C) that was previously described in another breast cancer cohort (Gaur et al., 2008). This AS event in *CEACAM1* affects 14 isoforms (Fig. 5B). The GMM clustering identified three tumor subpopulations (S1-S3) with distinct exon 7 inclusion levels, revealing that the exon can be variably included or skipped in the TCGA cohort (Fig. 5A). Exon 7 of *CEACAM1* was skipped in one of the breast cancer patients subpopulations, *CEACAM1*-S1, yet was preferentially included in normal breast from TCGA, normal breast tissue from GTEx as well as in several normal tissues such as lung, liver, heart, brain, blood and adipose tissue (Fig. 5B, Fig. S6A). The *CEACAM1*-S1 subpopulation with increased exon 7 skipping had worse overall survival when compared to the *CEACAM1*-S2 subpopulation (Fig. 5C), thus linking the SE event to an unfavorable disease outcome.

**Fig. 5.**
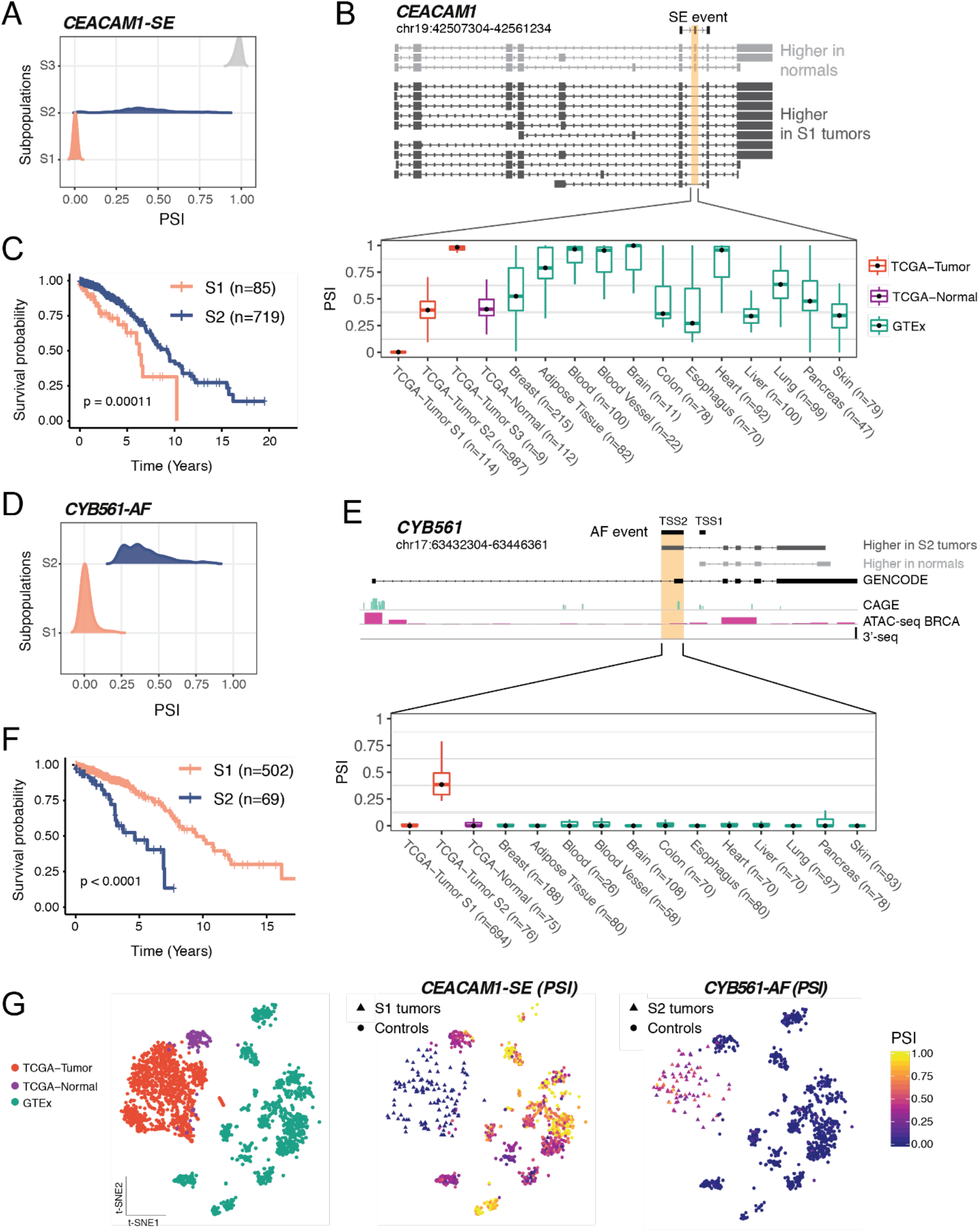
AS events in *CEACAM1* and *CYB561* are tumor-specific and associated with unfavorable prognosis in TCGA. **(A-C)** *CEACAM1* exon 7 skipping correlates with poorer overall survival in a subpopulation (S1) of breast cancer patients. TCGA tumor subpopulations (S1-S3) detected by Gaussian Mixture Models (GMM) clustering express different inclusion levels (percent spliced in or PSI) of exon 7 in *CEACAM1* (A). Structure of *CEACAM1* isoforms (gene coordinates are indicated) detected by LR-seq in breast tumors (dark grey) or normal tissues (light grey), highlighting the location of the skipped exon 7 (B, top panel). *CEACAM1* exon 7 inclusion (PSI) is shown in TCGA tumor subpopulations S1, S2, and S3, as well as in normal adjacent breast tissue from TCGA and a panel of GTEX normal tissues (B, bottom panel). Kaplan-Meier analysis of overall survival in TCGA breast cancer patients in the S1 subpopulation, which exhibits *CEACAM1* exon 7 skipping (n=85), and the S2 subpopulation, which exhibits higher exon 7 inclusion (n=719) (log-rank test, *P*<0.00011) (C). **(D-F)** Alternative first exon in *CYB561* (gene coordinates are indicated) correlates with poor overall survival in a subpopulation (S2) of breast cancer patients. TCGA subpopulations (S1-S2) detected by GMM clustering express different inclusion levels (PSI) of the alternative first exon in *CYB561* (D). Structure of *CYB561* isoforms detected by LR-seq in breast tumors (dark grey) or normal tissues (light grey), highlighting the location of novel (TSS1) or known alternative (TSS2, ENST00000581573) transcriptional start sites (E, top panel). CAGE, ATAC-seq and 3’-seq genomic tracks are displayed underneath isoform structures. Splicing of the isoform containing the *CYB561* TSS2 (as PSI) is shown in TCGA tumor subpopulations S1 and S2, as well as normal adjacent breast tissue from TCGA and a panel of GTEX normal tissues (E, bottom panel). Kaplan-Meier analysis of overall survival in TCGA breast cancer patients in the S1 subpopulation, which exhibits lower TSS2 inclusion (n=502), and the S2 subpopulation, which exhibits higher TSS2 inclusion (n=69) (log-rank test, *P*<0.0001) (F). (**G**) t-SNE representations of the AS events in *CEACAM1* and *CYB561.* Left, Samples are colored according to the dataset (Breast TCGA-tumor, Breast TCGA-Normal and GTEx). Right, Samples are colored according to the PSI levels of *CEACAM1*-SE and *CYB561*-AF events. The tumor subpopulations with differential splicing in each event are depicted in triangles, and controls are shown in circles.

Our analysis also identified a breast cancer-specific AF event involving two isoforms of *CYB561*, including a novel isoform identified by LR-seq. The GMM clustering detected two tumor subpopulations with distinct transcriptional start sites, TSS1 (novel) and TSS2 (known) (Fig. 5D-E). Patients in the *CYB561-*S2 subpopulation have a higher utilization of the isoform originating at the TSS2 start site, in comparison to the *CYB561-*S1 subpopulation and control tissues (Fig. 5E, Fig. S6B). Moreover, the *CYB561-*S2 subpopulation exhibited worse overall survival when compared to the *CYB561-*S1 subpopulation (Fig. 5F), thus linking the AF event to an unfavorable disease outcome. This AF event involves a novel isoform not annotated in GENCODE v.30, with a start site (TSS1) supported by CAGE and ATAC-seq peaks (Fig. 4B, Fig. 5E). The detection of this AF event was only possible due to the incorporation of this novel LR-seq isoform in the disease transcriptome. *CYB561* is an electron carrier enzyme that was recently identified as a novel prognostic factor in breast cancer (Shimizu and Nakayama, 2019).

We also identified a loss of intron retention in the breast cancer oncogene *E2F4* affecting 215 (19%) of TCGA breast tumors (Fig. 4B, Fig. S7). The GMM clustering identified three subpopulations with different splicing of *E2F4* isoforms in the TCGA cohort (Fig. S7A). In the *E2F4*-S1 subpopulation, *E2F4* switches from an intron-containing transcript in normal tissues that is not translated to a protein coding isoform in which the intron is spliced out in breast cancer patients (Fig. S7). The *E2F4* loss of intron retention is highly specific to breast cancer (Fig. 4B, Fig. S7). Interestingly, increased expression of the *E2F4* transcription factor is associated with cancer severity and poorer prognosis in breast cancer (Khaleel et al., 2014). In line with these findings, patients with loss of *E2F4* intron retention (*E2F4*-S1 subpopulation) have unfavorable prognosis when compared with patients where the intron is retained (Fig. S7). Thus, this retained intron event might represent a regulatory mechanism by which splicing leads to up-regulation of the *E2F4* oncogene in breast cancer.

In summary, patient stratification using our novel GMM clustering analysis identified tumor-specific splicing events in breast cancer that are confined to patient subpopulations with variable prevalence. These patient subpopulations carry distinctive splice alterations compared to tumor adjacent and normal tissues (Fig. 5G, Fig. S7E), and the differential splicing within these confined subpopulations could only be identified after patient stratification. We validate several of these novel isoforms in breast cancer cell lines (Fig. S8). Additionally, our analyses implicated genes regulated by AS such as *CEACAM1, CYB561* and *E2F4* as potentially playing a role in disease outcome.

## DISCUSSION

We performed LR-seq on 30 breast tumor and normal samples to define the full-length isoform-level transcriptome of human breast cancers, and developed an analytical pipeline to predict the functional consequences of cancer-associated splicing changes. We identified isoform-level diversity in tumors, and developed a thorough LR-seq-based breast cancer isoform catalog for quantitative and qualitative assessment of potential translation and subsequent protein domain effects. The data are made available through an interactive web portal which provides tools for querying, visualizing, and downloading data. We integrated LR-seq isoforms with orthogonal data sets to demonstrate the reliability of the approach, and proposed an analysis framework to detect the functional consequences of spliced isoforms in cancer. Finally, we used the resultant long-read breast cancer transcriptome to uncover novel isoforms associated with patient survival in TCGA using a Gaussian Mixture Model clustering approach to identify clusters of patients with similar splicing profiles.

Our pipeline uncovers 142,514 isoforms in breast tumors, 66% of which are novel when compared with the reference transcriptome, thereby significantly increasing the repertoire of known cancer isoforms. Although short-read RNA-seq adequately supports FSM isoforms, it is unable to detect our novel NIC and NNC isoforms at similar rates, pointing to the necessity of LR-seq to accurately define isoform-level transcriptomes. Many of our novel isoforms are supported by orthogonal data, such as CAGE and ATAC-seq for transcription start sites, and 3’-seq data for 3’ UTRs, supporting the validity of our findings. Until now, in breast cancer, LR-seq data were available on a small number of cell lines (Weirather et al., 2015; Nattestad et al., 2018; Lian et al., 2019), but not for primary tumors as described here. The proposed analysis likely captures the inter-tumor heterogeneity of primary tumor samples (Koren and Bentires-Alj, 2015), which is absent from cell lines, and provides a more clinically-relevant repertoire of spliced isoforms. This catalog of isoforms provides a more accurate and complete transcriptome enabling analyses at the isoform resolution in breast cancer, and possibly other cancer types. This long-read breast cancer transcriptome will likely help the discovery of novel targets for cancer therapeutics.

Even though several spliced isoforms for breast cancer genes such as *ESR1* and *ERBB2* have been previously identified (Thomas and Gustafsson, 2011; Volpi et al., 2019), current annotations widely used for transcriptome analysis, such as RefSeq and GENCODE, do not contain the level of complexity revealed by our LR-seq analysis. Importantly, a subset of the novel spliced isoforms contain distinct protein sequences, leading to novel combinations of protein domains and changes in cellular localization, and thus may play a role in promoting tumorigenesis or escaping drug response. For example, we uncovered a novel *ESR1* isoform predicted to gain a transmembrane domain and swap its localization from the nucleus to the cell membrane. Although point mutations as well as *ESR1* amplifications have been linked with breast cancer metastasis and therapy resistance (Lei et al., 2019), the role of *ESR1* splicing in the tumor response to endocrine therapies remains to be determined. A systematic characterization of isoform-level variation and complexity in tumors as described herein will help understand how isoforms might contribute to the heterogeneity of drug responses.

We identified AS events with prognostic value in TCGA breast cancer patients. Out of 310,861 AS events detected in our LR-seq transcriptome, we found 3,059 cancer-specific AS events, including 35 AS events were associated with significant changes in patient survival. Of these survival-associated events, 21 are absent from GENCODE and 10 are enriched in specific breast cancer subtypes. This analysis reveals that while AS events are frequent in cancer transcriptomes, they are mostly restricted to subpopulations of patients. However, several AS events are recurrent and affect more than half of TCGA patients, impacting: *UBAP2L*, a ubiquitin-associated protein upregulated in breast tumors and implicated in breast cancer cell cycle control (He et al., 2018)*; GPBP1*, a GC-Rich promoter-binding protein previously implicated in resistance to cisplatin and PARP inhibitors in ovarian cancer (Hu et al., 2018); and *CCDC120*, an interaction partner of the ADP-ribosylation factor 6 which is associated with breast cancer cell invasion (Hashimoto et al., 2004). The analysis also uncovered a novel regulatory mechanism by which the oncogenic E2F4 transcription factor is upregulated in breast tumors (Khaleel et al., 2014), linking an intron retention event in *E2F4* with unfavorable prognosis in breast cancer patients.

In conclusion, LR-seq is particularly well suited for the discovery of isoforms containing novel targets for immuno-oncology. These include the identification of cell surface isoforms against which specific monoclonal antibodies can be generated for use as therapeutics or as backbones for CAR-T cells. Isoforms also generate peptides which could be used for vaccination protocols, possibly in combination with checkpoint inhibitors.

## ACKNOWLEDGMENTS

We thank Drs. Charles Lee and Edison Liu for critically reading this manuscript and for their support in the development of this analytical platform; Dr. Taneli Helenius for scientific editing; Rahul Maurya, Jennifer Idol and Chew Yee Ngan and Genome Technologies Core at JAX-GM for help with LR-seq and RNAseq; members of the genomic core facility at the Icahn School of Medicine at Mount Sinai for help with LR-seq; PDX core at JAX-MG for providing samples; Research IT at JAX-GM for maintaining the website running the R/Shiny application; Prashant Singh, Francis O’Neill, Valeria Ochoa and members of the Anczuków lab for discussions. We also acknowledge the TCGA and GTEx consortiums for providing data analyzed in this study. This work was supported by start-up funds from the Jackson Laboratory to J.B. and O.A.; Director Innovation Funds award and a pilot project from the JAX NCI Cancer Center to J.B.; NIGMS grant R35GM133600 to C.R.B. Based on the data from this paper, JAX and Sanofi entered into a collaborative agreement. Part of D.F.T.V. and Y.Z. salaries were covered by this collaborative agreement.

## AUTHOR CONTRIBUTIONS

D.F.T.V. conceived and developed the methodology, performed bioinformatic analyses, and wrote the manuscript. A.N., Y.Z., and A.D.M. performed bioinformatic analyses. A.N., R.H., R.R., and T.W. performed experiments. K.P. provided reagents, expertise and feedback. O.A. and C.R.B advised in methodology development, provided expertise and wrote the manuscript. J.B. designed the study, acquired funding and revised the manuscript.

## MATERIALS AND METHODS

### Clinical samples

The study was conducted following approval by the Institutional Review Board (IRB) of The Jackson Laboratory for Genomic Medicine (IRB no. 16-NHSR-15, 17-JGM-06, 2018-039). Normal breast samples were acquired from the Maine Cancer Biospecimen Portal. Breast cancer tissue sections were contributed by Dr. Palucka. Exempt primary tissues from patients with breast cancer were obtained from the Baylor University Medical Center (BUMC) Tissue Bank (IRB no. 005-145; otherwise discarded tissues). Consecutive postsurgical tumor samples (from patients with in situ, invasive ductal, lobular, and/or mucinous carcinoma of the breast) were collected between years 2006 and 2013.

### PDX tumor samples lines

Snap frozen PDX tumor samples were obtained from The Jackson Laboratory (Sacramento, CA; catalog numbers provided in Table S1 as tumor identifiers). Upon receipt, frozen PDX tumors were placed in cryomolds (VWR, catalog# 4557), embedded in Optimal Cutting Temperature (O.C.T.) media (VWR, catalog# 4583) and stored at −80°C prior to RNA extraction.

### Human cell lines

Cell lines were purchased from the American Type Culture Collection (ATCC, Manassas, VA). CAMA-1, T-47D, and BT-549 lines were cultured in Dulbecco’s Modified Eagle’s Medium (ThermoFisher, catalog# 11965118) supplemented with 10% fetal bovine serum (Gemini Bio, catalog# 100-500). Hs578t was cultured in Dulbecco’s Modified Eagle’s Medium (ThermoFisher, catalog# 11965118) supplemented with 10% fetal bovine serum (Gemini Bio, catalog# 100-500) and 0.01mg/ml bovine insulin (Sigma Aldrich, catalog# I0516). Hs578Bst was cultured with Hybri-Care Medium (ATCC, catalog#. 46-X) and supplemented with 1.5 g/L sodium bicarbonate (ThermoFisher, catalog# 25080094), 30 ng/ml mouse EGF (ThermoFisher, catalog# PMG8043), and 10% fetal bovine serum (Gemini Bio, catalog# 100-500). MCF-7 was cultured in Eagle’s Minimum Essential Medium (ATCC, catalog# 30-2003) supplemented with 0.01 mg/ml recombinant human insulin and 10% fetal bovine serum. MDA-MB-231 and MDA-MB-468 lines were cultured in Leibovitz’s L-15 medium (ATCC, catalog# 30-2008), supplemented with 10% fetal bovine serum. HCC-1500 was cultured with RPMI-1640 Medium (ATCC, catalog# 30-2001) supplemented with 10% fetal bovine serum (Gemini Bio, catalog# 100-500). MCF-10A was cultured in MEBMTM Mammary Epithelial Cell Growth Basal Medium (Lonza, catalog# CC-3151) supplemented with 100ng/ml cholera toxin (Sigma Aldrich, catalog# C8052), BPE 2.00 mL (Lonza, #CC-3150), hEGF 0.50 mL (Lonza, #CC-3150), Insulin 0.50 mL (Lonza, #CC-3150), and Hydrocortisone 0.50 mL (Lonza, #CC-3150). Cell lines were kept at 37°C with 5% CO2.

### RNA extraction

High quality RNA was extracted from primary tumor tissues or cells. Briefly, using a Cryostat, four to five 0.3μm tissue sections were cut from OCT-embedded tumors and mixed in 350μ-Mercaptoethanol, and either frozen at −30°C or sent immediately for RNA extraction. For primary cells, cells were pelleted by centrifugation and then lysed with 350μl RLT + 10% β-Mercaptoethanol. RNA was extracted using the RNeasy Mini Prep Kit following the manufacturer’s instructions (Qiagen, catalog# 74106). Samples were treated with DNase I (Qiagen, catalog# 79254 and eluted in 30-50μl RNAse-Free water. RNA quality and quantity were assessed using a Qubit 2.0 fluorometer (Thermo Fisher Scientific), and only samples with RNA integrity number > 8.0 were selected for sequencing.

### Long-read library preparation and sequencing

Following RNA extraction, full-length cDNA synthesis from poly-A containing transcripts was performed using the Clontech SMARTer PCR kit (Takara). The resulting cDNA was PCR amplified to generate 1-2ug cDNA, and PCR products were purified using AMPure XP magnetic beads (Beckman Coulter). Size selection was performed using the Sage Science BluePippin System in order to remove small cDNA fragments which were preferentially sequenced by diffusion loading. Next, SMRTbell adapters were ligated to cDNA ends and purified by magnetic beads using the SMRTbell Template Prep Kit (Pacific Biosciences), followed by sequencing in a PacBio RSII or Sequel instrument. The list of of clinical samples, PDXs and cell lines sequenced in this study is provided in Table S1. Table S2 provides additional information including equipment, size selection and sequencing metrics for individual library runs.

### Short-read RNA sequencing

RNA was extracted using the Qiagen RNeasy Mini Prep kit and measured using a Qubit 2.0 Fluorometer (Thermo Fisher Scientific). RNA underwent quality control testing using a 2100 Bioanalyzer (RNA 6000 Pico kit, Agilent) followed by cDNA library preparation using the KAPA Stranded mRNA-Seq kit (Roche) according to manufacturer’s instruction. Paired end sequencing was performed at 76 base pairs on each side of the DNA fragment on the Illumina NextSeq platform. In total, RNA-seq was performed for 29 tumor and control samples, with 10.7-142.4 million reads sequenced per sample (mean = 46.4 million).

### PCR validation of AS events associated to survival

1ug RNA was reverse transcribed using the SuperScript IV First-Strand Synthesis System with both oligod(T) and random hexamer primers per manufacturer instructions (Invitrogen, #18091050). Touhc-down PCR was used to amplify 200ng of cDNA with Q5 High-Fidelity DNA Polymerase and the High-GC content buffer (New england Biolabs, #M0491L) and primers listed in Supplementary Table S4 on a BioRad T100 Thermal Cycler (BioRad, #1861096). PCR products were separated in 2% agarose gel stained with SYBR Safe (Invitrogen) and imaged using ChemiDoc MP Imaging System (Bio-rad).

### Long-read data processing

Raw PacBio Iso-seq data (BAM files) were processed using the ToFU pipeline (Gordon et al., 2015) obtained from https://github.com/PacificBiosciences/IsoSeq_SA3nUP/wiki. Briefly, the pipeline generates non-redundant full-length (FL) transcripts in the following steps: (i) classify reads as FL reads and non-FL reads based on the presence of adapters and polyA signal, (ii) identify isoform clusters for each transcript using FL read(s), (iii) polish isoform sequences by performing error correction and obtaining a final consensus transcript using both FL and non-FL reads. FL transcripts were mapped to human (hg38) and mouse (mm10) genomes using gmap (Wu and Watanabe, 2005), and transcripts aligned to mouse were discarded from downstream analyses (PDX samples). FL transcripts for all samples were merged using chain_samples.py from the cDNA_Cupcake tools (https://github.com/Magdoll/cDNA_Cupcake) to create a non-redundant merged transcriptome. In addition to the samples sequenced in this study, a publicly available MCF7 cell line dataset (Anvar et al., 2018) was re-processed and included in the merged transcriptome (see Table S2).

### RNA-seq data processing

Fastq files were aligned to the hg38 genome using hisat2 v. 2.0.4 (Pertea et al., 2016) with default options, followed by removal of duplicate reads with samtools v. 1.3.1. Bigwig files were generated using bamCoverage v.3.3.0 from deeptools2 (Ramírez et al., 2016). Xenome v.1.0 (Conway et al., 2012) was used to filter out mouse and ambiguous reads in PDX samples. External RNA-seq datasets were retrieved from the dbGAP database using the following accession numbers: The Cancer Genome Atlas (TCGA, phs000178.v11.p8) and Genotype-Tissue Expression (GTEx; phs000424.v8.p2).

### Isoform annotation and quality control

Our isoform annotation pipeline combined several tools for isoform transcript and open-reading frame annotation as outlined in Figure S3A. Spliced isoforms were annotated with SQANTI (Tardaguila et al., 2018), using GENCODE comprehensive v.30 as reference. We also used SQANTI2 (https://github.com/Magdoll/SQANTI2) to obtain a comprehensive set of quality attributes for sequenced FL reads at both transcript and junction level, that were applied for retaining high-quality transcripts and filtering out potential artifacts as detailed below. First, SQANTI was used to generated an indel-corrected FASTA/GTF files by realignment of FL transcripts to the human genome hg38, as well as to classify isoforms based on their splice patterns using GENCODE v.30 as reference and to flag transcripts with non-canonical junctions or junctions possibly derived from reverse-transcriptase switching (RT-switching). SQANTI2 was used to compute junction coverage in the Intropolis dataset and distance of TSS to CAGE peaks. Gffcompare (Pertea et al., 2016) was used to annotate transcripts with potential intron retention (class codes *m*, *n*, *i* and *y*). In general, novel isoforms (ISM, NIC, NNC) were filtered based on 3’end reliability (no poly(A) intra-priming), non-canonical junctions or RT-switching junctions and splice junction support as described below.

The read coverage filter applied to novel transcripts was defined as follows: the transcript was kept if all splice junctions were covered by at least 5 short-reads (RNA-seq from Intropolis dataset) or the transcript was detected in at least 3 Iso-seq samples (i.e., minimum of 3 full-length reads). Additionally, to remove potential poly-A intra-priming during the reverse transcriptase reaction, the genomic 3’end of a transcript was considered unreliable if it had all the following properties: (i) it was located more than 100bp away from an annotated TTS, (ii) the adenine percentage downstream of TTS > 80%, (iii) and no overlap with the polyAsite database (Gruber et al., 2016), a catalog of high-quality and curated poly(A) sites detected by 3’seq. Based on the combination of these quality attributes, we devised the following filtering strategy for each transcript category:

FSM: no filtering (all included)
ISM: filtering out transcripts with unreliable 3’ends
NIC: filters based on unreliable 3’ends, minimum read coverage, and no intron-retention
NNC: filters based on unreliable 3’ends, minimum read coverage, no intron-retention, no junctions labeled as RT-switch, and only canonical splice sites

Other (intergenic, antisense, fusion of adjacent loci): minimum read coverage, no intron-retention, no junctions labeled as RT-switch, and only canonical splice sites

### Protein-level functional characterization of long-read isoforms

We used sequence homology and domain conservation to human protein isoforms in UniProt to determine optimal coding sequences from full-length LR-seq isoforms as described below. First, we assembled a comprehensive human proteome reference including both canonical (SwissProt+TrEmbl) and spliced isoforms (VarSplice) from UniProt release 2019-04, which contained 95,915 protein sequences. Possible coding sequences (ORFs) from LR-seq isoforms were predicted using Transdecoder (Haas et al., 2013), and local alignment using blastp (Camacho et al., 2009) was performed against the reference proteome using options max_target_seqs = 1 and e-value = 10^-5^ to identify homologs in UniProt. Also, PFAM domains for all extracted ORFs were predicted using the hmmscan tool from hmmer v.3.1 (http://hmmer.org) using default parameters. Then, a single best ORF for each transcript was selected based on significant sequence homology (blastp) and domain conservation (hmmer) to human proteins.

Next, we performed extensive annotation of coding sequences using multiple tools and custom scripts. Prediction of transmembrane helices was carried out using TMHMM, and subcellular localization was inferred using DeepLoc. Global alignment of FL coding sequences to homologs in UniProt was carried out using the Needleman-Wunsch algorithm implemented in the pairwiseAlignment function from the Biostrings package in R. Nonsense-mediated mRNA decay (NMD) analysis was performed using a custom R script using the coding sequence predicted by Transdecoder. Specifically, a FL transcript was predicted as NMD sensitive when the stop codon occurred before the terminal exon and was located more than 55 nucleotides upstream of the last splice junction. Scripts for performing global alignment and NMD prediction were implemented using mclapply (parallel package v.3.4.1).

### Peptide search

Raw MS/MS datasets from TCGA breast cancer patients (230 samples from 125 tumors) were retrieved from the CPTAC database (Mertins et al., 2016). In addition, 45 breast cancer samples from another patient cohort (Johansson et al., 2019) were obtained from the ProteomeXchange database, for a total of 275 proteomic samples. Peptide identification was performed using MS-GF+ version 2018.10.15 (Kim and Pevzner, 2014) using a sequence database that contained 165,477 ORFs derived from long-read isoforms, in addition to 95,915 human protein sequences from UniProt, and 116 contaminant sequences. The following parameters were set for database searching: Carbamidomethyl (C), iTRAQ4plex (N-term), and iTRAQ4plex (K) were specified as fixed modifications. Oxidation (M), Deamidated (NQ), Acetyl (K), and Methyl (K) were specified as variable modifications. The precursor mass tolerance for protein identification on MS was 20 ppm, and the product ion tolerance for MS/MS was 0.05 Da. Partial cleavage by trypsin was used, with up to two missed cleavages permitted. MZID profiles identified from the search engine were then pooled using the R/Bioconductor package MSnbase (Gatto and Lilley, 2012), and peptide-to-spectrum matches (PSM) satisfying both spectra and peptide FDRs cutoffs < 1% were kept for further analysis. Finally, PSMs were classified into four types, namely unique_PacBio (peptides uniquely mapped to a single FL isoform and not mapped to UniProt), non_unique_PacBio (peptides mapped to more than one FL isoform and not mapped to UniProt), non-unique-PacBio+Uniprot (peptides mapped to both PacBio and Uniprot proteins), and multigene (peptides mapped to multiple genes).

### Splice event extraction and quantification in TCGA and GTEx samples

Alternative splicing events were extracted and quantified using SUPPA2 (Trincado et al., 2018), based on a GTF containing long-read isoforms merged to GENCODE v.30. The input transcript expression file containing TPM abundances of all isoforms for SUPPA2 was computed using StringTie v.1.3.0 (Pertea et al., 2016) using the merged GTF as a reference for transcript quantification. Alternative first and last exons were defined using 250 base pairs overlap threshold (*-t* 250) for PSI calculation. For other types of events (SE, MX, RI, A3, A5) the overlap threshold was set to 10 base pairs.

### Gaussian mixture clustering and survival analysis

The Gaussian mixture models (GMM) clustering was implemented in R using the mclust package v. 5.4.1. The clustering approach consisted in fitting a mixture of Gaussian distributions to PSI values from AS events simultaneously in cancer and control samples, using the PSI matrix obtained from SUPPA2. Model fitting with mclust was performed using one to three Gaussian distributions (*i.e.*, minimum of one and maximum of three PSI subpopulations), and the optimal fitting was determined using the Bayesian information criterion. For each AS event, samples were assigned to clusters (subpopulations) with highest probability, and the frequency of tumor and controls were computed within subpopulations. Subpopulations with high tumor purity (>90% breast tumor samples) were further analyzed for differential splicing and survival as described below. The Wilcox ranksum test in R was used to determine differential splicing between a tumor-specific subpopulation and control tissues from TCGA and GTEx. Next, survival analysis was done using the pairwise_survdiff function from the survminer package v. 0.4.6, which performs pairwise comparisons between GMM-inferred subpopulations with corrections for multiple testing. Only subpopulations with at least 30 patients were included in the Kaplan-Meier analysis. Significant survival events were selected based on the global p-value and pairwise comparisons (adjusted P < 0.01).

### t-SNE visualization of AS events

t-distributed stochastic neighbor embedding (t-SNE) (Maaten and Hinton, 2008) was performed using the Rtsne package v 0.13. The t-SNE representation was generated based on the PSI matrix of exon skipping events. The PSI matrix was filtered to remove events with more than 80% of missing values. Samples with more than 80% of missing values were also removed. Missing values occurred for events in which neither the inclusion or skipping forms are detected. Any remaining missing values were mean imputed. t-SNE with learning rate 200 and perplexity 50 were applied for all visualizations.

## Supplemental Information

**Fig. S1.**
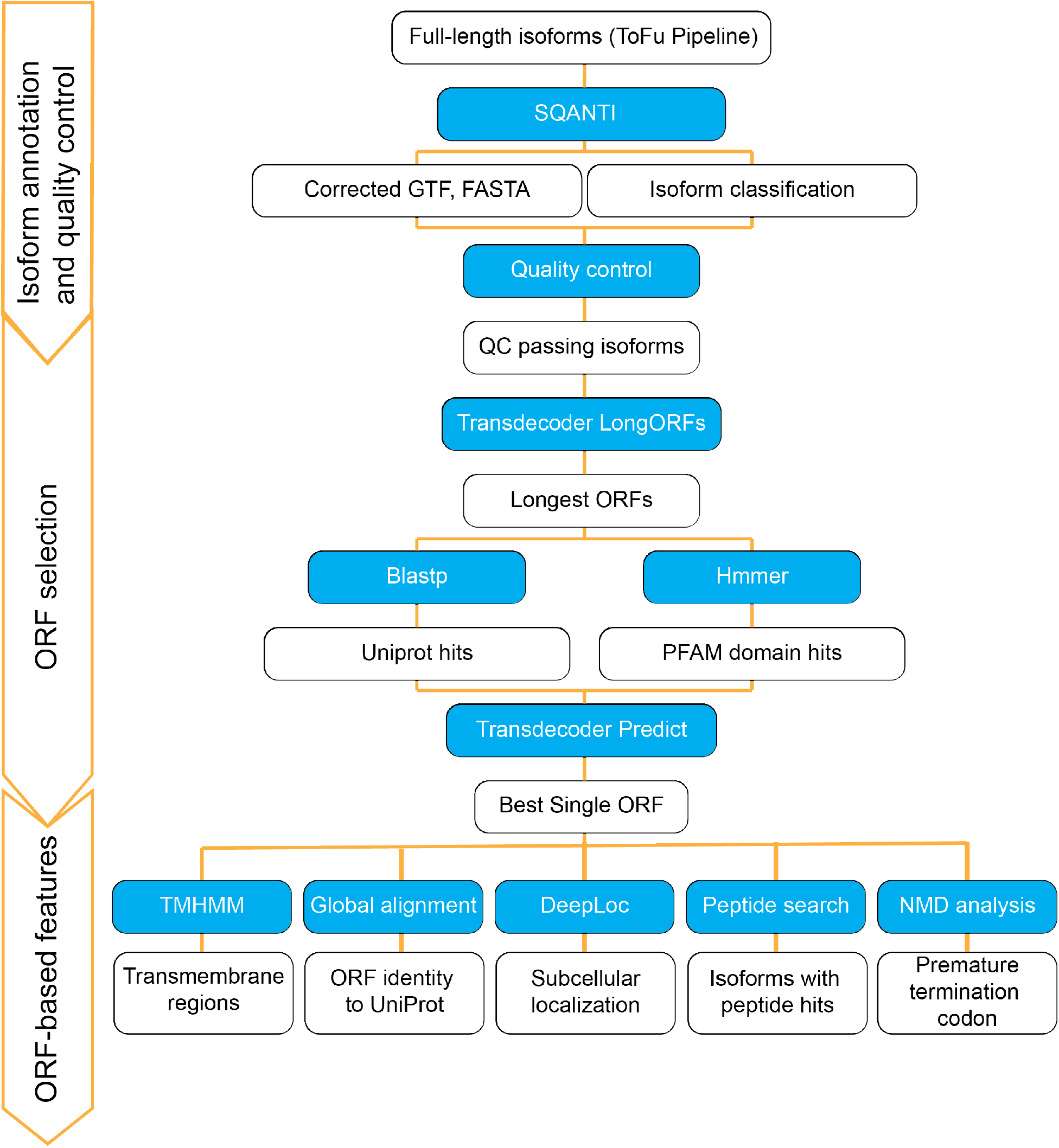
Analysis pipeline for LR-seq isoform annotation, quality control and prediction of protein features (related to Fig. 1). The pipeline is divided in three phases: 1) isoform annotation and quality control, 2) ORF selection, and 3) prediction of ORF-based protein features. In the first phase, SQANTI and SQANTI2 are used to obtain quality metrics for LR-seq isoforms at both transcript and junction levels, which are applied for removing low quality long-reads containing inadequate splice junction support, and those that contained signatures of poly(A) intra-priming or non-canonical junctions derived from reverse transcriptase template switching. Second, the optimal ORF selection is performed using a combination of Transdecoder, blastp and hmmer tools. The third phase consists of downstream analysis such as global alignment of ORFs to UniProt, inference of non-mediated decay (NMD), peptide search in proteomics datasets, and prediction of protein features using TMHMM and DeepLoc. Blue boxes refer to tools and scripts used for data processing, while white boxes refer to inputs and outputs. See Methods for additional details.

**Fig. S2.**
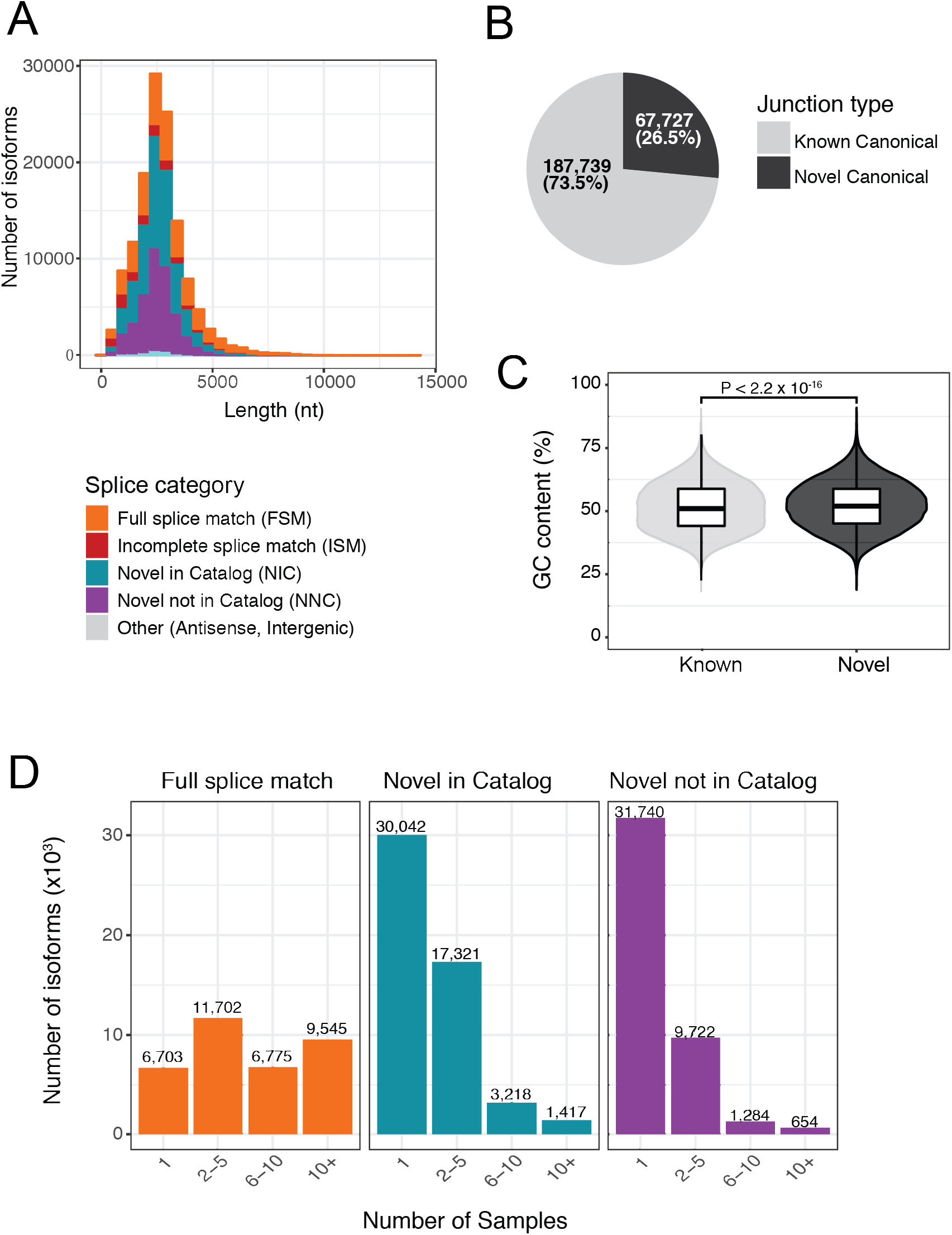
Characteristics of isoforms and splice junctions detected by LR-seq (related to Fig. 1). **(A)** Length distribution (nt) of LR-seq isoforms detected, colored by structural category as described in A. **(B)** Frequency and absolute number of novel and known unique splice junctions detected in the LR-seq breast cancer transcriptome. **(C)** GC content of known or novel canonical splice junctions detected in the LR-seq breast cancer transcriptome in a region −50bp to +50bp from the splice site. The 2% increase in the mean GC content surrounding novel junctions is significant by a Wilcox rank sum test (*P* < 2.2 x 10^-16^). **(D)** Frequency of detection of LR-seq isoforms across libraries, colored by structural category as described in A.

**Fig. S3.**
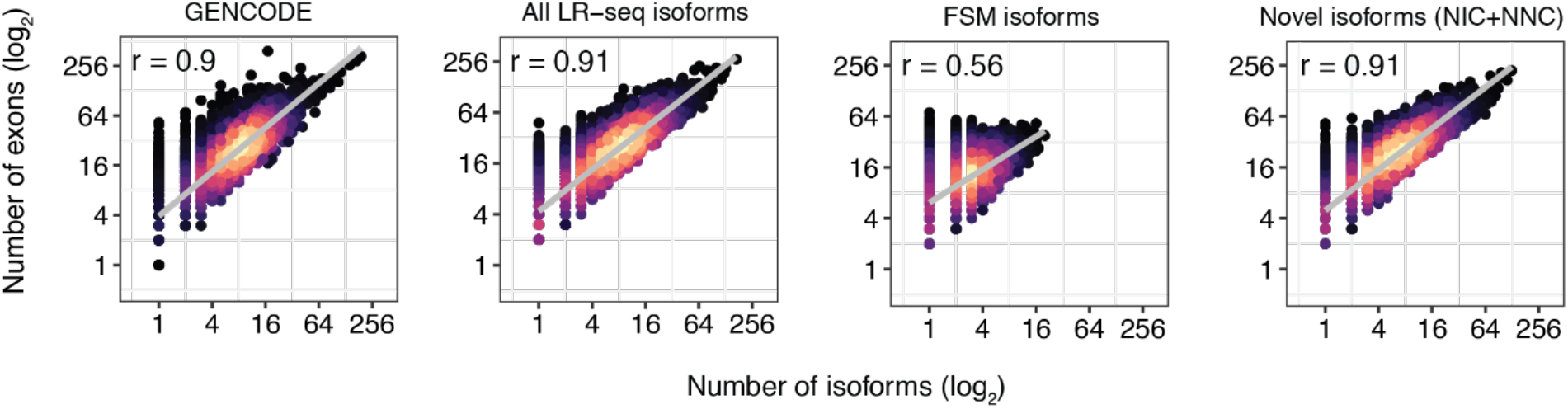
Correlation between number of exons and number of LR-seq isoforms (related to Fig. 1). From left to right: isoforms in GENCODE v.30, all LR-seq isoforms detected in this study, FSM isoforms only, and novel isoforms only (NIC and NNC). Spearman correlations are indicated. See Fig.1A for FSM, NIC and NNC definitions.

**Fig. S4.**
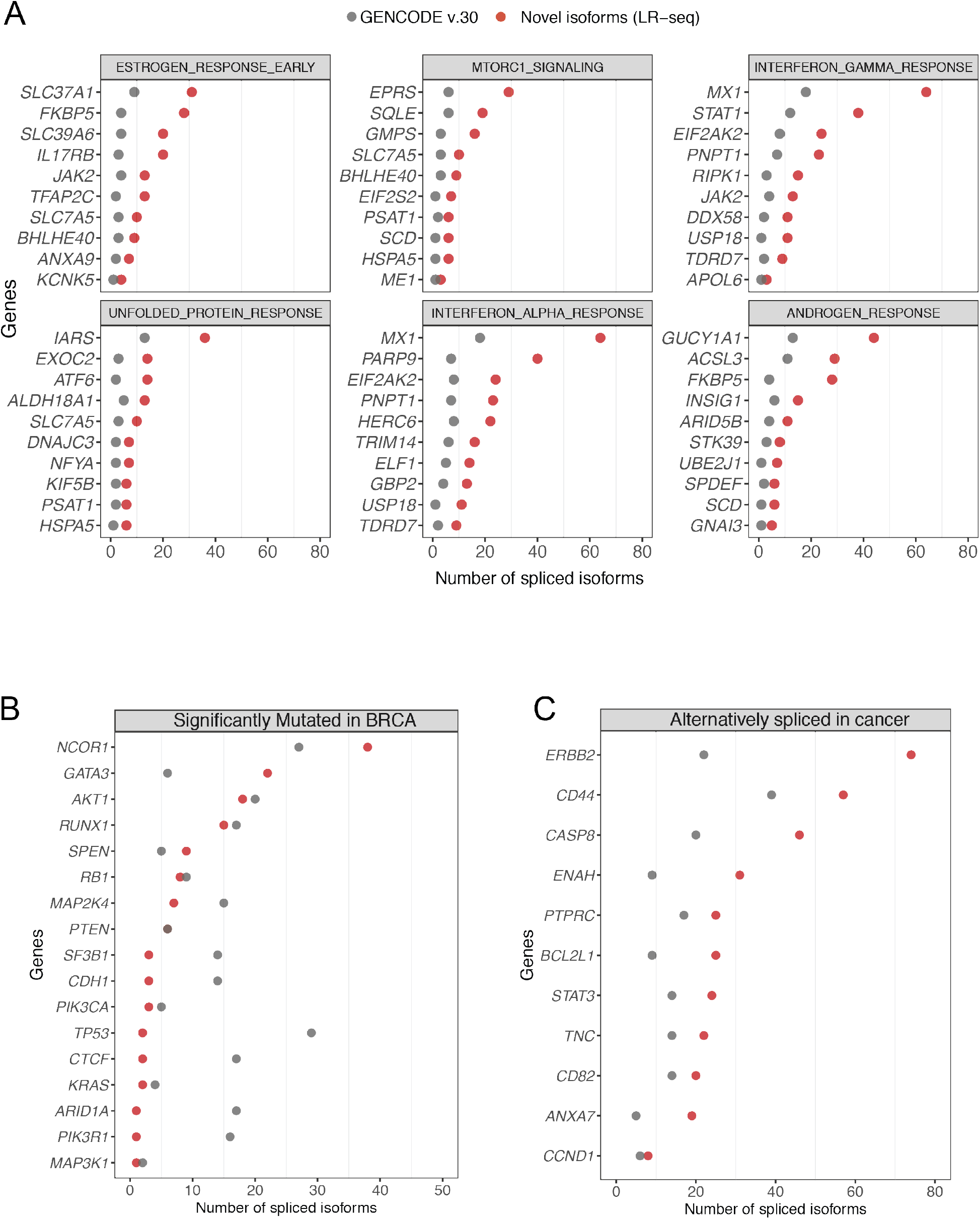
Isoform increase in cancer-associated genes and pathways (related to Fig. 2). **(A)** The top ten genes with the highest number of novel LR-seq isoforms are shown for each pathway from Fig. 2B compared to known isoforms from GENCODE v.30. **(B)** Number of novel isoforms detected by LR-seq *vs.*GENCODE v.30 in significantly mutated breast cancer genes (http://www.tumorportal.org). **(C)** Number of novel isoforms detected by LR-seq *vs.*GENCODE v.30 in genes that have been previously reported to undergo AS in cancer (Sveen et al., 2016; Urbanski et al., 2018).

**Fig. S5.**
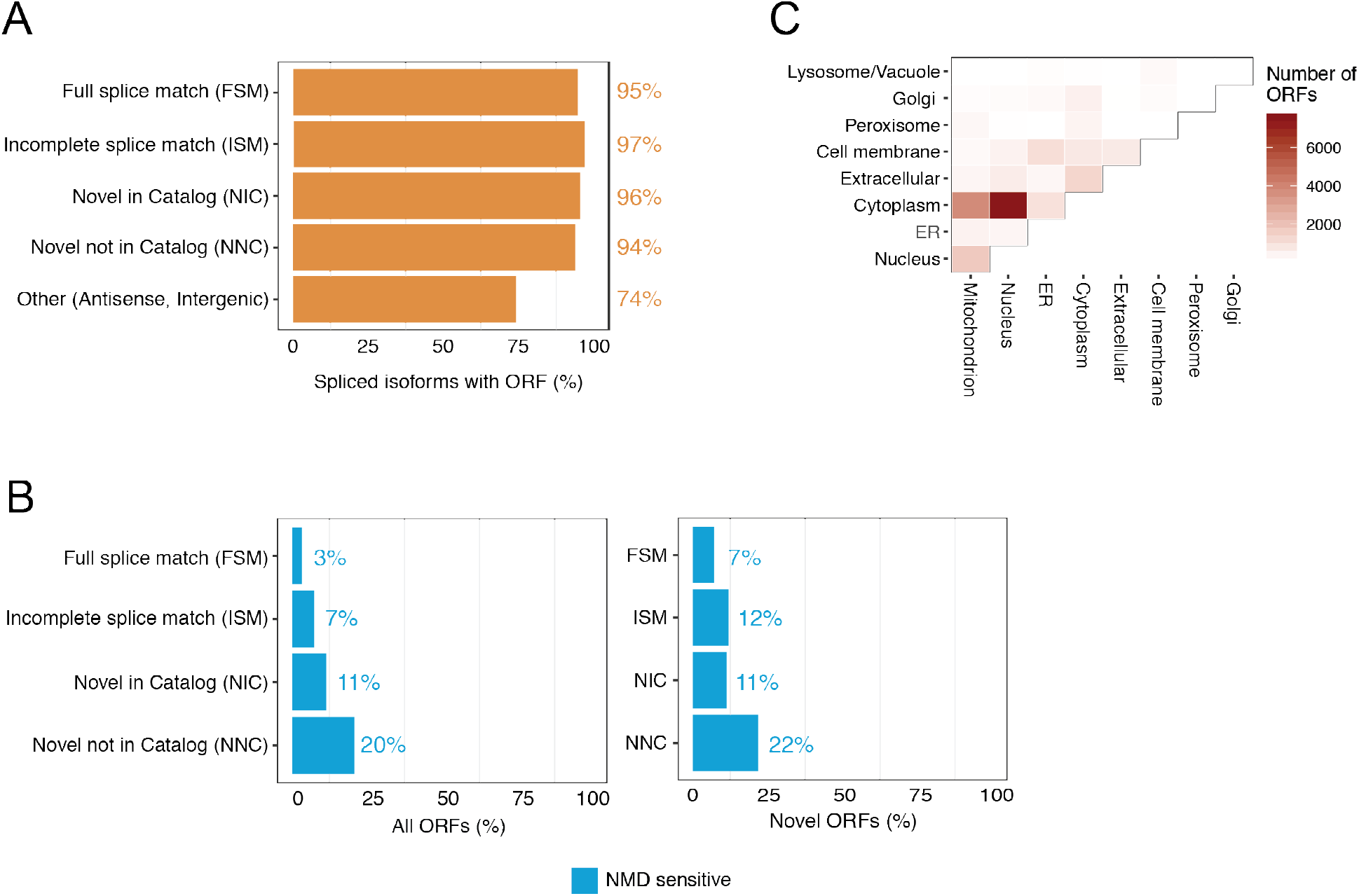
Coding potential and NMD prediction of ORFs extracted from LR-seq isoforms detected in breast tumors (related to Fig. 3). **(A)** Percent of LR-seq isoforms predicted by Transdecoder to contain an ORF, plotted per isoform structural category from Fig. 1A. **(B)** Percent of LR-seq isoforms classified as NMD sensitive (containing a premature termination codon before the last exon) per isoform structural category for all ORFs or from novel-only ORFs. Novel ORFs have < 99% sequence similarity to their closest human protein variant in UniProt. **(C)** DeepLoc-predicted changes in cellular localization for ORFs derived from LR-seq isoforms, compared to their closest human protein isoform in UniProt. The scale indicates the number of LR-seq isoforms changing their localization between each cellular localization pair.

**Figure S6.**
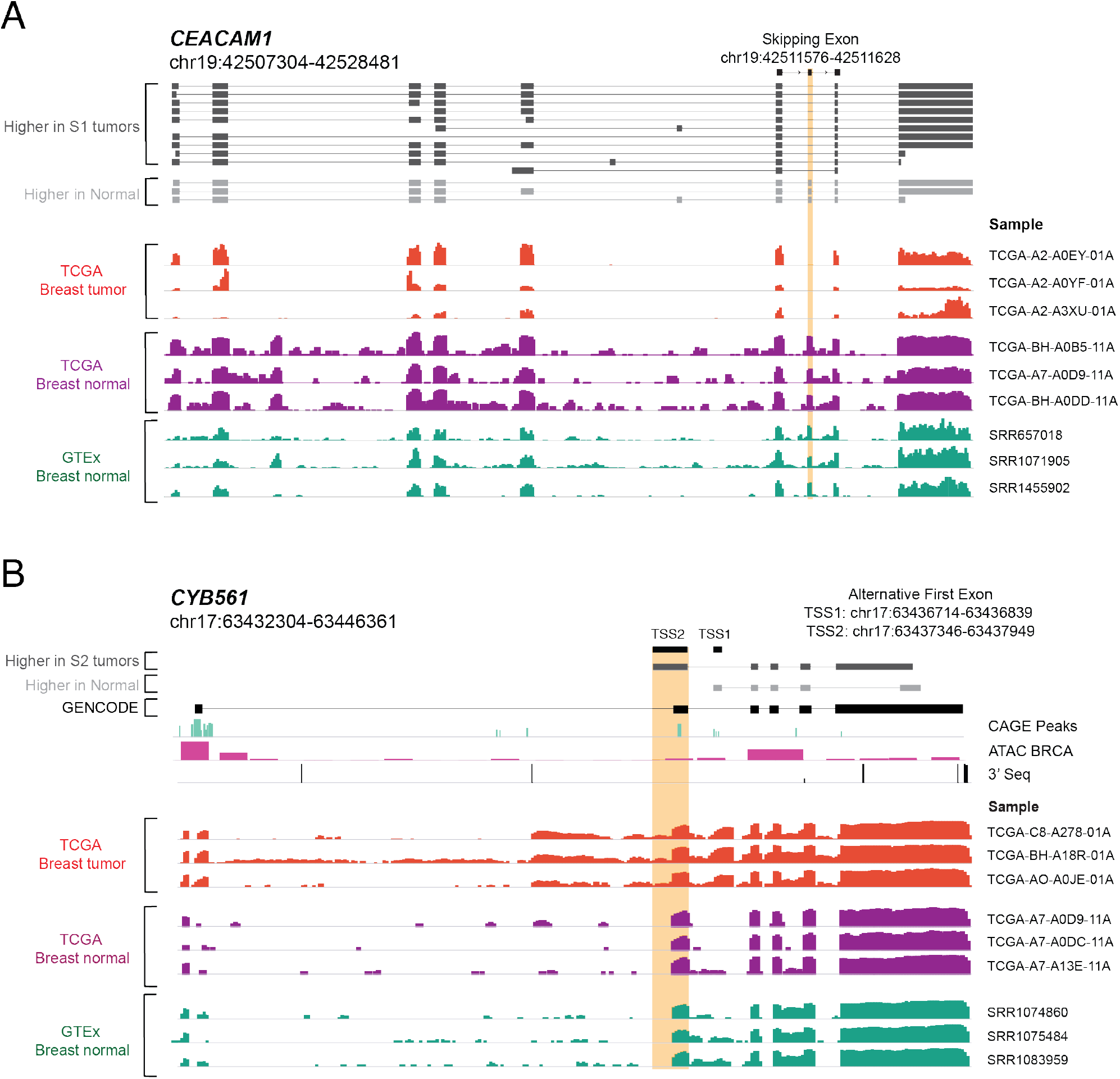
RNA-seq coverage plots of AS events in *CYB561* and *CEACAM1* in tumors and normal tissues (related to Fig. 5). **(A)** RNA-seq coverage tracks from the *CEACAM1* isoform associated with skipped exon 7 (gene coordinates are indicated) are shown for selected TCGA (n=3), as well as normal breast tissue from TCGA (n=3) and from GTEx (n=3). Structure of *CEACAM1* isoforms detected by LR-seq in breast tumors (dark grey) or normal tissues (light grey), highlighting the location and genomic coordinates of the skipped exon 7. TCGA and GTEX sample identification numbers are listed next to each track. **(B)** RNA-seq coverage tracks from the *CYB561* isoform associated with an alternative first exon (gene coordinates are indicated) are shown for selected TCGA breast tumors (n=3), as well as normal breast tissue from TCGA (n=3) and from GTEx (n=3). Structure of *CYB561* isoforms detected by LR-seq in breast tumors (dark grey) or normal tissues (light grey), highlighting the location and genomic coordinates of known (TSS1) and novel (TSS2) transcriptional start sites. CAGE, ATAC-seq and 3’-seq genomic tracks are displayed underneath isoform structures. TCGA and GTEX sample identification numbers are listed next to each track.

**Figure S7.**
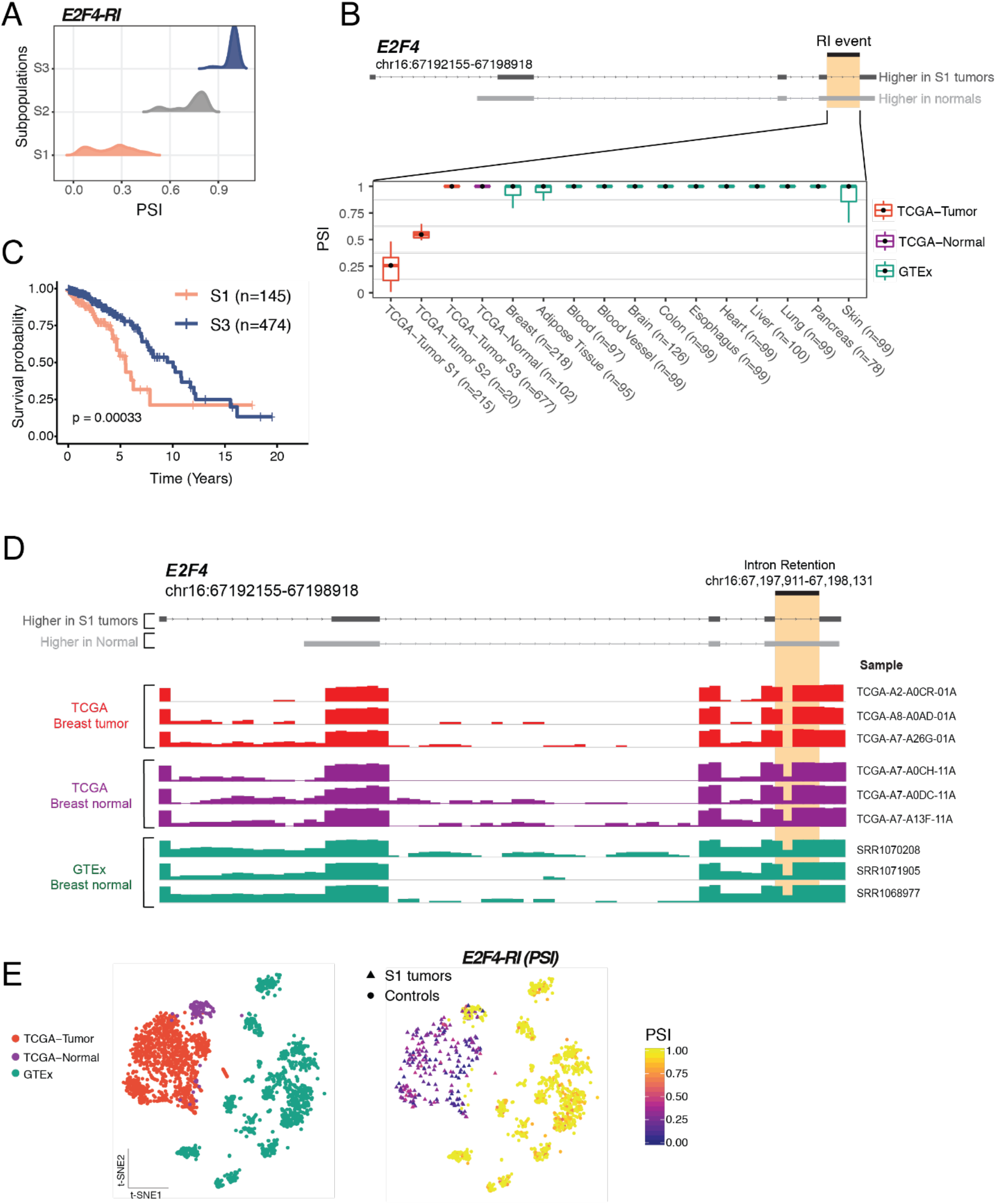
Loss of intron retention in *E2F4* in a tumor subpopulation correlates with unfavorable prognosis (related to Figs. 4 and 5). **(A)** TCGA subpopulations (S1-S3) detected by GMM clustering express different inclusion levels (percent spliced in or PSI) of an intronic region in *E2F4*. **(B)** Structure of *E2F4* isoforms detected by LR-seq in breast tumors (dark grey) or normal tissues (light grey), highlighting the location of the retained intron (B, top panel). *E2F4* intron retention (as PSI) is shown in TCGA tumor subpopulations S1, S2, and S3, as well as in normal adjacent breast tissue from TCGA and a panel of GTEX normal tissues (B, bottom panel). **(C)** Kaplan-Meier analysis of overall survival in TCGA breast cancer patients in the S1 subpopulation, which exhibits lower intron retention (n=145), and the S3 subpopulation, which exhibits higher intron retention (n=474) (log-rank test, *P*<0.0003) (C). **(D)** RNA-seq coverage tracks from the *E2F4* isoform associated with a retained intron event (gene coordinates are indicated) are shown for selected TCGA breast tumors (n=3), as well as normal breast tissue from TCGA (n=3) and from GTEx (n=3). Structure of *E2F4* isoforms detected by LR-seq in breast tumors (dark grey) or normal tissues (light grey), highlighting the location and genomic coordinates of retained intron. TCGA and GTEX sample identification numbers are listed next to each track. **(E)** t-SNE representations of the AS event in *E2F4.* Left, Samples are colored according to the dataset (Breast TCGA-tumor, Breast TCGA-Normal and GTEx). Right, Samples are colored according to the PSI levels of intron retention in *E2F4*. Tumor subpopulation with differential splicing is depicted in triangles, and controls are shown in circles.

**Figure S8.**
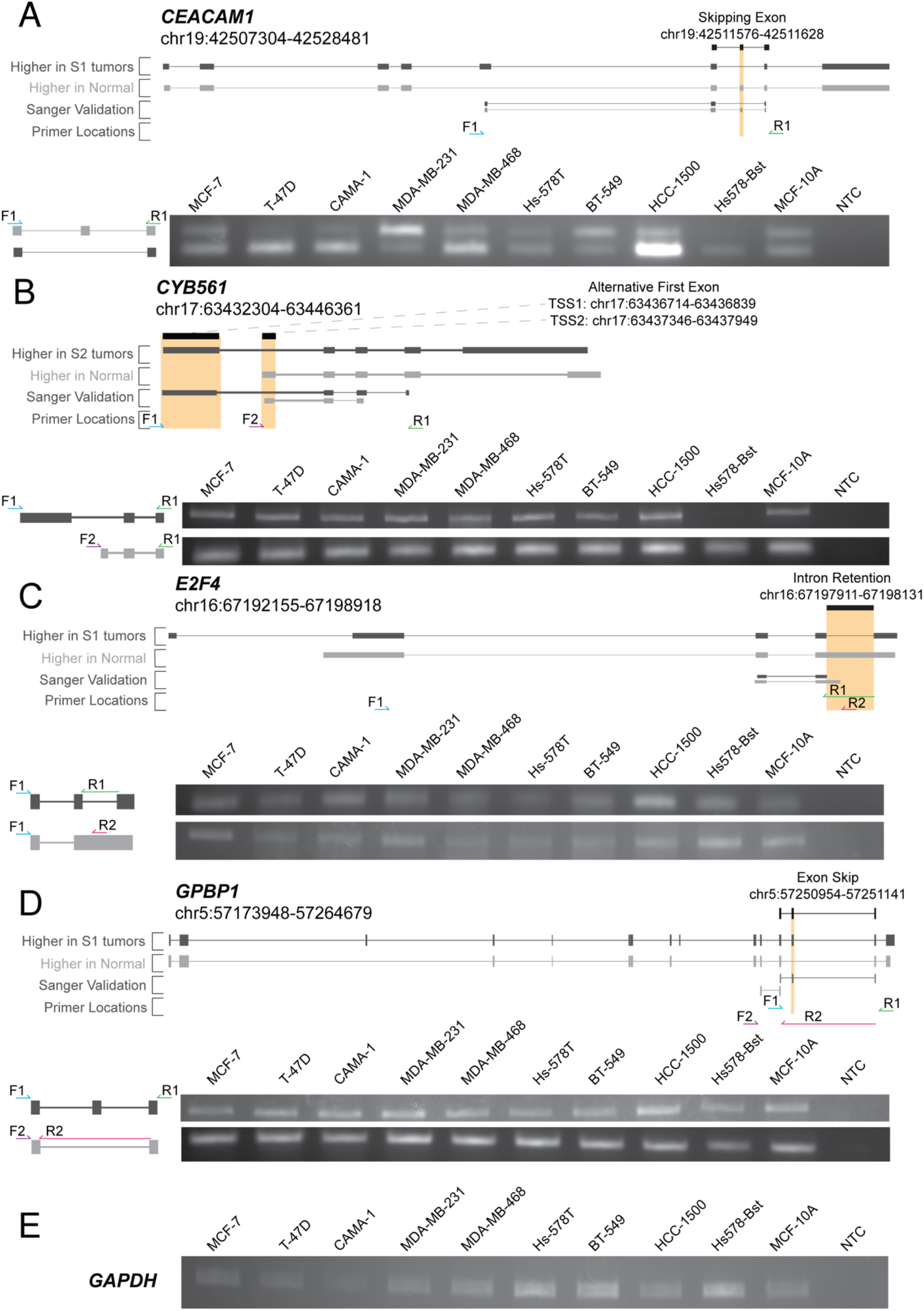
Detection in breast cancer cell lines of the tumor-specific AS events associated with unfavorable prognosis identified in TCGA breast tumors (related to Figs. 4 and 5). **(A-D)** RT-PCR detection of AS events in *CEACAM1* (A), *CYB561* (B), *E2F4* (C), or *GPBP1* (D), in breast cancer (MCF-7, T-47D, CAMA-1, MDA-MB-231, MDA-MB-468, Hs-578T, BT-549, HCC-1500) or non-transformed mammary epithelial (Hs578-Bst, MCF-10A) cell lines using isoform specific primers that amplify both the included and skipped isoforms detected by SUPPA in TCGA breast tumors. For each event, the gene coordinates, the structure of isoforms detected by LR-seq in breast tumors (dark grey) or normal tissues (light grey), highlighting the type, location and coordinates of the AS event are indicated in the top panel; an orange box highlights the differentially spliced sequence. The sanger validation tracks show the alignment of the amplified product for a pooled set of breast cancer cell lines. The primer locations track depicts that PCR primers positions. A representative gel is shown along with the isoform structures and primer locations. NTC-no template control. **(E)** RT-PCR detection of housekeeping GAPDH transcript as a loading control.

**Table S1.** Histology and demographics of breast cancer clinical samples, PDXs and cell lines profiled by LR-seq in this study.

**Table S2.** Sequencing metrics, equipment, and size selection parameters for individual PacBio library runs.

**Table S3.** Alternative splice events and associated isoforms with survival correlation in TCGA breast cancer patients.

**Table S4.** Primers for PCR validation of AS events in *CYB561*, *CEACAM1*, *E2F4*, and *GPBP1*.

